# O-Acyltransferase Genes Involved in the Production of Volatile Sex Pheromones in *Caenorhabditis elegans*

**DOI:** 10.1101/2025.08.10.669515

**Authors:** Xuan Wan, Sarah M Cohen, Yan Yu, Henry Hoan Le, Heenam Park, Alessandro Groaz, Rachel Moreno, Minyi Tan, Matthew R. Gronquist, Ryoji Shinya, Frank C. Schroeder, Paul W Sternberg

## Abstract

Gene family expansions are critical for functional diversification, yet paralog contributions to metabolic pathways are often unclear. In *Caenorhabditis*, the expanded O-acyltransferase (OAC) family—enzymes that transfer acyl groups to hydroxylated substrates—remains poorly characterized despite having been implicated in lipid metabolism. Using CRISPR-Cas9 mutagenesis, behavioral assays, gas chromatographic-mass spectral (GC-MS) analyses, and metabolomics, we systematically analyzed 59 OAC-family protein-coding genes to define their roles in regulating signaling molecules. We found that four adjacent paralogs (*oac-13, oac-16, oac-25,* and *oac-28*) on chromosome I are required for synthesizing volatile sex pheromones (VSPs)—airborne signals critical for male mate-searching. Specifically, oac*-13* and *oac-16* are necessary for producing both major pheromone components, while the identical tandem paralogs *oac-25* and *oac-28* regulate the production of the later-eluting component in gas chromatography. Disruption of these genes reduced production of key pheromone components and impaired male attraction. Metabolomics revealed that *oac-16* and other OACs also modulate synthesis and secretion of non-volatile ascaroside pheromones, indicating dual roles in chemical signaling. This work uncovers functional specialization within an expanded gene family, illustrating how redundancy and divergence enable adaptive evolution of communication systems.

## Introduction

The nematode *Caenorhabditis elegans* has served as an important model organism for decades, owing to its transparency, compact genome, and fully mapped neural connectome, features that enable unparalleled genetic and neurobiological studies ^1–3^. *C. elegans* has approximately 20,000 genes ^2,4^. Only about 4000 have any described phenotype or other information derived from nematode-based *in-vivo* or *in-vitro* experiments, excluding bulk gene expression measurements and RNAi information ^5^. About half the genes have names, indicating either due to phenotypes directly studied in *C. elegans,* for homology with researched genes in other model organisms or to established protein domains ^6^. By comparison, about 40% of human genes have some known functions, and two-thirds of this information comes from model organisms ^7^. Thus, systematic analyses are important for understanding *C. elegans* especially given its role as a model for nematodes in general and for human genes.

O-acyltransferases are a subset of the acyltransferase class of enzymes that catalyze the transfer of acyl groups onto hydroxyl groups. This contrasts them with N-acyltransferases which transfer acyl groups onto amines. O-acyltransferases are ubiquitous throughout the cellular environment and are used in pathways ranging from bacterial growth of biofilms, to Wnt signaling in flies, to the conversion of cholesterol to cholesteryl ester in humans ^8–10^. The most commonly studied forms of O-acyltransferases are the membrane-bound O-acyltransferases, which include ACAT1 and PORCN and their orthologs in *Drosophila* and other model organisms ^11,12^. *C. elegans* has several families of membrane-bound O-acyltransferases, such as *mboa* (membrane-bound O-acyltransferase) family, *mom* (more of MS), and the *oac* (O-acyltransferase) family ^5^. The *mboa* gene family (10 genes), and the *mom* gene family (5 genes) all contain a membrane-bound O-acyltransferase domain (MBOAT) (InterPro IPR004299; Pfam PF03062); and are related to genes in the Wnt and Hedgehog pathways ^9,13^. However, the largest yet least-studied class of O-acyltransferases in *C. elegans* is named OAC gene family, which contains a different protein domain, the acyltransferase 3 domain (InterPro IPR002656; Pfam PF01757) (Figure 1b). The OAC class of enzymes was named due to its predicted ability to transfer acyl groups other than amino-acyl groups, but it has otherwise been unstudied in *C. elegans*.

**Figure 1.**
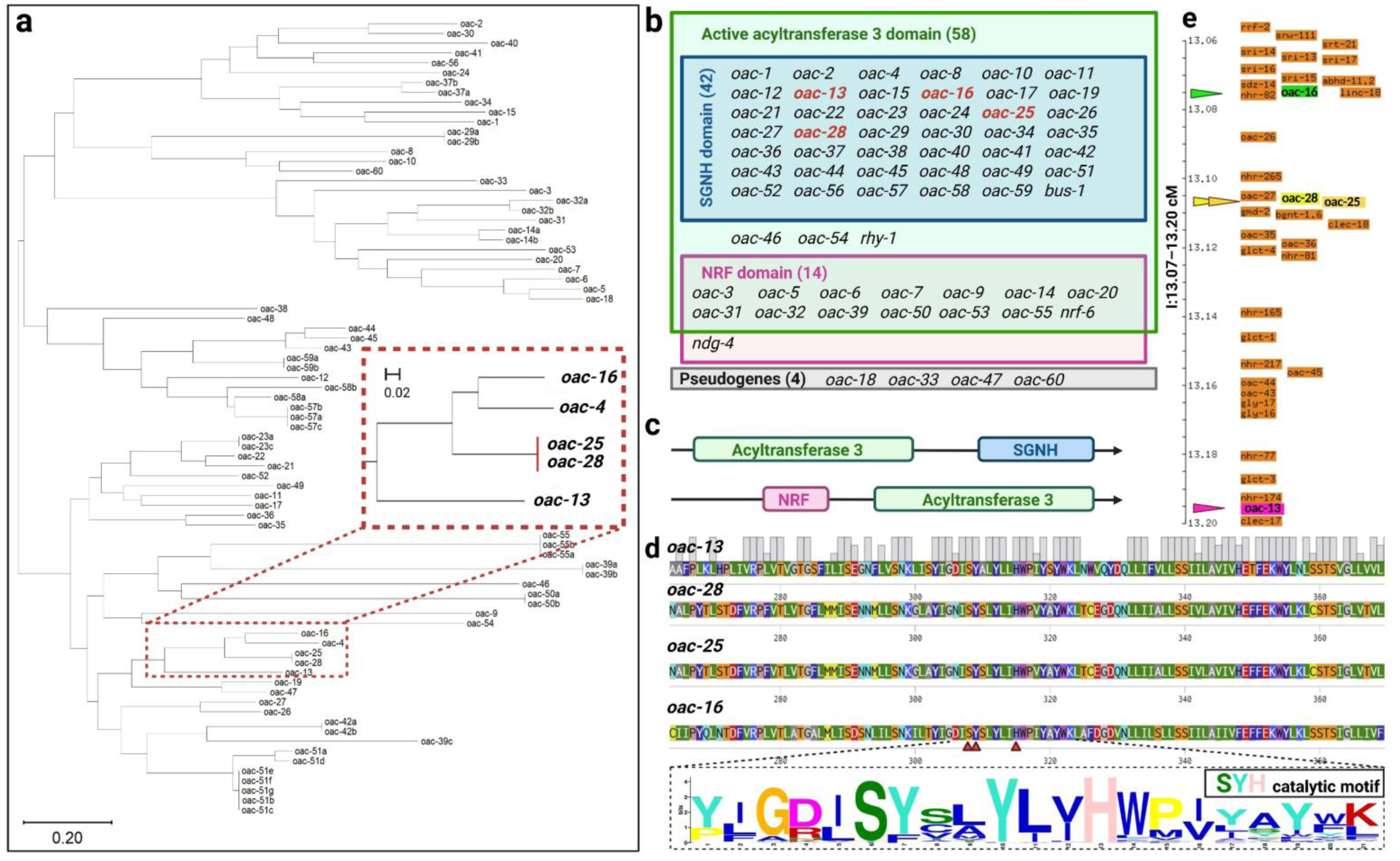
The OAC gene family is defined by an acyltransferase 3 domain. **(a)** Phylogenetic tree of OAC proteins in *C. elegans*. The length of branches indicates the evolutionary distance between proteins. **(b)** The genes of the broader OAC gene family are separated by name and protein domain architecture. Typically, proteins encoded by *oac* genes have two protein architectures: **(c)** an acyltransferase 3 domain with a C-terminal SGNH hydrolase-type esterase domain or an acyltransferase 3 domain with a N-terminal nose resistant-to-fluoxetine (NRF) domain. **(d)** Top panel: conserved SYH catalytic motif in OAC acyltransferase 3 domains. Multiple sequence alignment of OAC proteins highlights the universally conserved Ser (S), Tyr (Y), and His (H) residues (indicated in red arrows). Bottom panel: the sequence logo of aligned acyltransferase 3 domains from 59 OAC genes (56 *oac* named genes, and *bus-1, nrf-6, rhy-1*). Letter height reflects residue conservation among those 59 genes (taller = higher conservation). **(e)** Genomic loci of VSPs related OAC family genes in *C. elegans*. *oac-13* (genomic position: Chromosome I: 12,421,239–12,424,411); *oac-16* (genomic position: Chromosome I: 12,239,160–12,241,808); *oac-25* (genomic position: Chromosome I:12269255..12272190); *oac-28* (genomic position: Chromosome I: 12259912..12262848) are located in close proximity on chromosome I.

In addition, the acyltransferase 3 domain is highly conserved across species. In bacteria, this domain facilitates cell surface modifications that confer resistance to lysosomes and bacteriophages ^14,15^. OAC is an evolutionarily ancient and conserved gene family, spanning prokaryotes (archaea, bacteria) and eukaryotes (fungi, nematodes, insects, mammals). Notably, insects and nematodes exhibit multi-copy retention of OAC genes in their genomes, mammals retain only a single orthologous copy ^16^. A nonsense mutation in acyltransferase 3 domain caused a loss of function of ACYL3 in humans and chimpanzees during the evolution of the great apes ^16^. *C. elegans* genome harbors 59 protein-coding *oac* genes ^5^.

Chemical communication is vital for *C. elegans* reproduction. The nematode employs two pheromone classes: non-volatile ascaroside pheromones, which regulate development and social behaviors ^17–22^, and VSPs that guide long-range mate navigation ^23–27^. Ascarosides comprise a chemical language within nematodes, regulating many aspects of development and social behaviors ^17–19,21,22,28,29^. The VSPs are produced exclusively by self-sperm/sperm-depleted hermaphrodites/females to attract males. Their structures and biosynthetic pathway remains elusive. Many studies have shown that OACs are essential to volatile insect pheromone production ^30–32^. The conservation of OAC-driven pheromone biosynthesis in other species raises compelling questions: Could *C. elegans*’s OAC family genes similarly regulate pheromone signaling? Why has the OAC family undergone expansion in *C. elegans*? Does this reflect functional redundancy, sub-functionalization, or novel adaptations unique to nematode ecology?

To fully allow us to understand the *C. elegans* organism, we need to systematically create mutants and examine gene families rather than relying on a shotgun approach. In this paper, we chose the under-studied OAC gene family to create strains and study their potential functions. In this study, we generated 56 mutants spanning the entire family and performed behavioral assays, GC-MS analysis, and comparative metabolomic profiling to assess their roles in pheromone production.

## Results

### Phylogenetic analysis of the OAC gene/protein family

We compared the protein sequences and created a protein phylogenetic tree (Figure 1a) to understand how the genes/proteins were related. We also found that four additional genes should be included in the OAC gene family (Figure 1b). Although there are several strains available that contain an *oac* mutation as well as numerous other mutations, these mutant strains were less useful in identifying the specific functions of OAC genes ^33^. Therefore, we created at least one isogenic mutant strain for each protein-coding *oac* for which there was not already an existing complete loss-of-function mutant ^34,35^. We studied the volatile sex pheromone production metabolomics of the OAC gene class as the first phenotypic analyses of these mutants.

The OAC family is defined by the acyltransferase 3 domain, a conserved catalytic motif essential for acyl group transfer. In addition to the named *oac* genes, we identified three additional genes including this OAC-specific acyltransferase-3 domain: *bus-1, rhy-1,* and *nrf-6*. These genes were named prior to the designation of the OAC gene class. These genes were previously characterized for roles unrelated to acyltransferase activity. *bus-1* is a gene that is expressed in the rectal epithelial cells, and *bus-1* mutants are unaffected by the nematode rectal pathogen *Microbacterium nematophilum*, which causes tail-swelling, hence the name <u>b</u>acterially <u>u</u>n<u>s</u>wollen ^36^. *rhy-1* is a regulator of the hypoxia-inducible factor HIF-1 through the *egl-9* pathway; mutants of *rhy-1* exhibit some egg-laying defects ^37^. Fluoxetine (Prozac) causes *C. elegans* nose-muscle contractions, yet nose resistant-to-fluoxetine (*nrf*) mutants are immune to this phenomenon ^38^. *nrf-6* mutants also display a pale-egg phenotype caused by a lack of yolk granules; this causes subsequent retarded development and partial embryonic lethality ^38^. Based on their shared domain architecture, we include *bus-1, rhy-1* and *nrf-6* in the OAC gene family. Following the inclusion of these genes, the OAC family comprises 64 genes, with four pseudogenes (*oac-18, oac-33, oac-47,* and *oac-60*) identified among them.

All 59 OAC protein-coding genes have an acyltransferase 3 domain. OACs are generally categorized into two groups based on their non-acyltransferase 3 domain architecture (Figure 1c): 42 OACs have an acyltransferase 3 domain coupled with a C-terminal SGNH hydrolase domain (InterPro IPR043968), while 15 OACs have an acyltransferase 3 domain with an N-terminal nose resistant-to-fluoxetine (NRF) domain (InterPro IPR006621) (Figure 1b). The SGNH hydrolase domain, often associated with carbohydrate modification ^15^, and the NRF domain suggest diverse functional roles for these enzymes. In general, membrane-bound O-acyltransferases have conserved histidine residues required for functional enzymatic activity ^39,40^. We find that all the OACs that have an acyltransferase 3 domain likewise have a conserved catalytic histidine residue and the combination of Ser (S)-Tyr (Y)-His (H) residues in the acyltransferase 3 domain as shown in the motif logo (SYH catalytic motif, Figure 1d, Supplemental Figure 1). Tandemly duplicated genes, such as *oac-21*/*oac-22*/*oac-23*; *oac-13/oac-16; oac-25/oac-28* tend to cluster together, suggesting recent gene duplication events. However, the overall phylogenetic structure does not correlate strongly with genomic loci, indicating functional divergence among paralogs.

### Strain construction for functional analysis

To systematically investigate the OAC gene family, we obtained or generated knockout strains for all 59 protein-coding *oac* genes. Existing mutants for *bus-1(e2678)*, *nrf-6(sa525)*, and *rhy-1(ok1402)* were obtained from the *Caenorhabditis* Genetics Center (CGC). For the remaining 56 genes, we employed CRISPR/Cas9-mediated genome editing to create 95 mutant alleles (Table 1; Supplemental Tables 1 and 2). We used two strategies: (1) non-homologous end joining to generate deletions of several hundred base pairs, and (2) homology-directed repair to introduce multiple early stop codons or frameshifts. These strains provide a comprehensive resource for probing the functional roles of OACs in *C. elegans*.

**Table 1.** 95 alleles representing 56 *oac* genes were made using either a deletion or knock-in CRISPR/Cas9 method.

| Genotype | Strain | Description |
| --- | --- | --- |
| *oac-1(sy1399)* | PS8547 | 605 bp Deletion |
| *oac-2(sy1218)* | PS8201 | Triple-Stop Knock-In |
| *oac-2(sy1219)* | PS8202 | Triple-Stop Knock-In |
| *oac-3(sy1329)* | PS7536 | 387 bp Deletion |
| *oac-3(sy1053)* | PS7577 | 1,428 bp Deletion |
| *oac-4(sy1400)* | PS8548 | 5 bp Deletion |
| *oac-5(sy1333)* | PS7556 | 854 bp Deletion |
| *oac-5(sy1334)* | PS7791 | 1,231 bp Deletion |
| *oac-6(sy1326)* | PS7466 | 1,899 bp Deletion |
| *oac-7(sy1327)* | PS7490 | 546 bp Deletion |
| *oac-8(sy1284)* | PS8365 | Triple-Stop Knock-In |
| *oac-8(sy1285)* | PS8366 | Triple-Stop Knock-In |
| *oac-9(sy1324)* | PS7592 | 1,129 bp Deletion |
| *oac-10(sy1339)* | PS7908 | 875 bp Deletion |
| *oac-10(sy1340)* | PS8110 | 942 bp Deletion |
| *oac-11(sy1407)* | PS8310 | 1438 bp Deletion |
| *oac-12(sy1378)* | PS8439 | Triple-Stop Knock-In |
| *oac-12(sy1379)* | PS8440 | Triple-Stop Knock-In |
| *oac-13(sy1367)* | PS8386 | 708 bp Deletion |
| *oac-13(sy1368)* | PS8387 | 481 bp Deletion |
| *oac-14(sy1343)* | PS7593 | 544 bp Deletion |
| *oac-14(sy1344)* | PS7696 | 868 bp Deletion |
| *oac-14(sy1345)* | PS7697 | 404 bp Deletion |
| *oac-15(sy1369)* | PS8388 | 579 bp Deletion |
| *oac-15(sy1370)* | PS8389 | 835 bp Deletion |
| *oac-16(sy1341)* | PS8113 | 1,255 bp Deletion |
| *oac-16(sy1342)* | PS8108 | 746 bp Deletion |
| *oac-16(sy1471)* | PS8577 | Triple-Stop Knock-In |
| *oac-17(sy1405)* | PS8299 | 419 bp Deletion |
| *oac-17(sy1406)* | PS8300 | 1,056 bp Deletion |
| *oac-19(sy1741)* | PS9361 | Triple-Stop Knock-In |
| *oac-20(sy1346)* | PS7950 | 757 bp Deletion |
| *oac-20(sy1347)* | PS7956 | 561 bp Deletion |
| *oac-20(sy1348)* | PS8109 | 988 bp Deletion |
| *oac-21(sy1566)* | PS8897 | Triple-Stop Knock-In |
| *oac-22(sy1286)* | PS8367 | Triple-Stop Knock-In |
| *oac-22(sy1287)* | PS8368 | Triple-Stop Knock-In |
| *oac-23(sy1384)* | PS8517 | Triple-Stop Knock-In |
| *oac-23(sy1385)* | PS8518 | Triple-Stop Knock-In |
| *oac-24(sy1246)* | PS8250 | Triple-Stop Knock-In |
| *oac-24(sy1247)* | PS8251 | Triple-Stop Knock-In |
| *oac-25(sy1620) oac-28(sy1621)* | PS9143 | 8 bp insert; 8 bp insert |
| *oac-26(sy1386)* | PS8519 | Triple-Stop Knock-In |
| *oac-27(sy1475)* | PS8714 | Triple-Stop Knock-In |
| *oac-27(sy1476)* | PS8715 | Triple-Stop Knock-In |
| *oac-28(sy1734)* | PS9351 | 8 bp insert |
| *oac-29(sy1335)* | PS7787 | 896 bp Deletion |
| *oac-30(sy1577)* | PS8934 | Triple-Stop Knock-In |
| *oac-30(sy1578)* | PS8935 | Triple-Stop Knock-In |
| *oac-31(sy1328)* | PS7559 | 591 bp Deletion |
| *oac-32(sy1330)* | PS7558 | 1,764 bp Deletion |
| *oac-34(sy1290)* | PS8394 | Triple-Stop Knock-In |
| *oac-34(sy1291)* | PS8395 | Triple-Stop Knock-In |
| *oac-35(sy1390)* | PS8538 | Triple-Stop Knock-In |
| *oac-35(sy1391)* | PS8539 | Triple-Stop Knock-In |
| *oac-36(sy1411)* | PS8580 | Triple-Stop Knock-In |
| *oac-36(sy1412)* | PS8581 | Triple-Stop Knock-In |
| *oac-37(sy1401)* | PS8549 | Triple-Stop Knock-In |
| *oac-38(sy1256)* | PS8265 | Triple-Stop Knock-In |
| *oac-38(sy1257)* | PS8266 | Triple-Stop Knock-In |
| *oac-39(sy1321)* | PS7428 | 546 bp Deletion |
| *oac-40(sy1258)* | PS8301 | Triple-Stop Knock-In |
| *oac-40(sy1259)* | PS8302 | Triple-Stop Knock-In |
| *oac-41(sy1413)* | PS8582 | Triple-Stop Knock-In |
| *oac-41(sy1414)* | PS8583 | Triple-Stop Knock-In |
| *oac-42(sy1433)* | PS8632 | Triple-Stop Knock-In |
| *oac-42(sy1434)* | PS8633 | Triple-Stop Knock-In |
| *oac-43(sy1381)* | PS8480 | Triple-Stop Knock-In |
| *oac-44(sy1431)* | PS8630 | Triple-Stop Knock-In |
| *oac-44(sy1432)* | PS8631 | Triple-Stop Knock-In |
| *oac-45(sy1742)* | PS9362 | Triple-Stop Knock-In |
| *oac-46(sy1747)/ mln1[dpy-10(e128) mls14]* | PS9372 | Triple-Stop Knock-In, balanced |
| *oac-46(sy1747)* | PS9373 | Triple-Stop Knock-In |
| *oac-48(sy1402)* | PS8550 | Triple-Stop Knock-In |
| *oac-49(sy1380)* | PS8479 | Triple-Stop Knock-In |
| *oac-50(sy1336)* | PS7716 | 559 bp Deletion |
| *oac-50(sy1337)* | PS7790 | 927 bp Deletion |
| *oac-50(sy1338)* | PS7730 | 782 bp Deletion |
| *oac-51(sy1358)* | PS8482 | Triple-Stop Knock-In |
| *oac-51(sy1359)* | PS8483 | Triple-Stop Knock-In |
| *oac-51(sy1388)* | PS8536 | Triple-Stop Knock-In |
| *oac-51(sy1389)* | PS8537 | Triple-Stop Knock-In |
| *oac-52(sy1288)* | PS8369 | Triple-Stop Knock-In |
| *oac-52(sy1289)* | PS8370 | Triple-Stop Knock-In |
| *oac-53(sy1331)* | PS7652 | 483 bp Deletion |
| *oac-53(sy1332)* | PS7687 | 592 bp Deletion |
| *oac-54(sy1322)* | PS7465 | 1,138 bp Deletion |
| *oac-55(sy1323)* | PS7717 | 1,030 bp Deletion |
| *oac-56(sy1372)* | PS8527 | Triple-Stop Knock-In |
| *oac-56(sy1373)* | PS8528 | Triple-Stop Knock-In |
| *oac-57(sy1374)* | PS8529 | Triple-Stop Knock-In |
| *oac-57(sy1375)* | PS8530 | Triple-Stop Knock-In |
| *oac-58(sy1376)* | PS8531 | Triple-Stop Knock-In |
| *oac-58(sy1377)* | PS8532 | Triple-Stop Knock-In |
| *oac-59(sy1260)* | PS8303 | Triple-Stop Knock-In |
| *oac-59(sy1261)* | PS8304 | Triple-Stop Knock-In |

None of the *oac* mutant strains exhibited observable behavioral phenotypes, except *oac-16(sy1342).* Worms homozygous for this allele displayed a locomotion defect, characterized by restricted movement away from the hatching site. However, the other two *oac-16* alleles (*sy1341, sy1471*) lacked this mobility defect, suggesting the *sy1342* phenotype stems from an off-target mutation, rather than *oac-16* loss of function. As this defect is unrelated to pheromone signals, we include *oac-16(sy1342)* strain in the pheromone extraction for GC-MS analysis.

### OAC-13, OAC-16, OAC-25, and OAC-28 are required to maintain the male-attracting feature of hermaphrodite-derived extracts

Given the established roles of OACs in lipid modification and pheromone biosynthesis in other species ^30–32^, we hypothesized that *C. elegans* OACs might similarly regulate VSPs production. Using behavioral assays (Figure 2) and GC-MS analyses (Figures 3 and 4), we identified four OACs required for VSPs biosynthesis.

**Figure 2.**
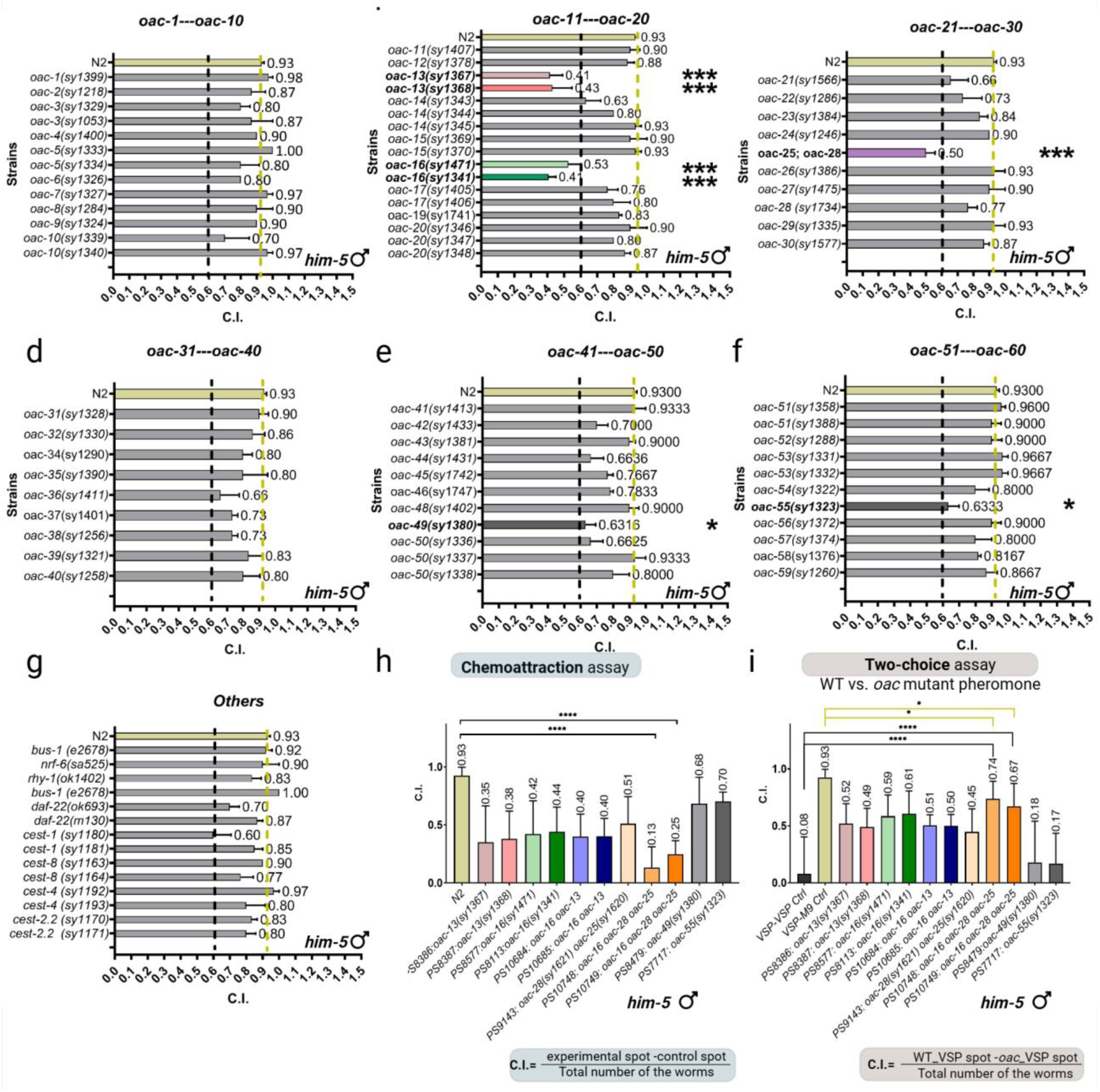
Comparative analysis of chemotactic responses to *C. elegans* VSPs. **(a-g)** Chemoattraction index of volatile pheromone extracts from wild type (N2) strain and 59 *oac* mutants, assayed with *him-5* males (See Methods). Most mutants retained wild type bioactivity, but *oac-13, oac-16, oac-28 oac-25* (double mutant of identical paralogs), *oac-49,* and *oac-55* exhibited significant reductions (below 0.6. Sample sizes: n = 60 males per trial, 3 trials. **(h)** Validation of VSPs signal defects in *oac-13, oac-16, oac-28 oac-25* (double mutant of identical paralogs), *oac-49,* and *oac-55* mutants in large-scale trials (n = 400 males per trial, 20 trials). **(i)** Two-choice preference assays showing male preference for wild type hermaphrodite extracts over mutant extracts (n = 400 males per trial, 20 trials). Mutants of tandem paralog pair *oac-13, oac-16, oac-16 oac-13* and *oac-28 oac-25* mutant exhibited reduced attractiveness compared to wild type extracts in both (h) and (i). The *oac-16 oac-13* double mutant showed no additive defects, indicating functional redundancy. Error bars: S.D.; P < 0.005; P < 0.05 (two-tailed unpaired t-test); ns., not significant (P > 0.05). The yellow dashed line represents the wild type N2 strain chemoattraction index. The black dashed line represents the screening threshold C.I.<0.6.

**Figure 3.**
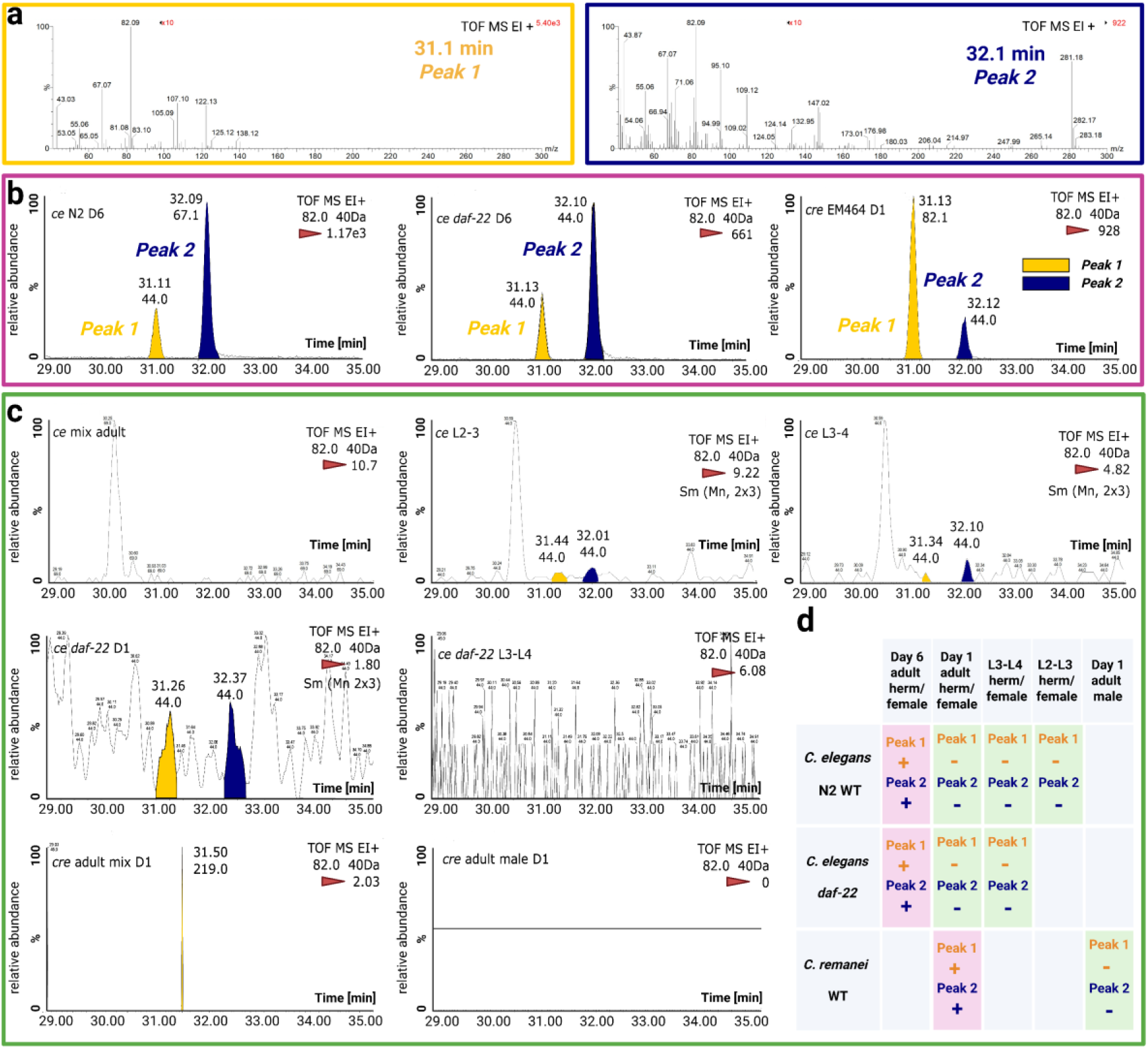
GC-MS analysis identifies two conserved chromatographic peaks associated with reproductive-stage-regulated VSPs production. **(a)** Representative mass spectra of *Peak 1* (t_R_=31.1 min) and *peak 2* (t_R_=32.1 min) detected in pheromone-positive samples. Retention times and mass spectra were conserved across samples. **(b)** Representative GC-MS extracted ion chromatograms (EIC) for m/z 82.0 of a pheromone-positive sample (6-day-old self-sperm-depleted *C. elegans* N2 and *daf-22* mutant hermaphrodites and 1-day-old not mated *C. remanei* female) showing *Peaks 1* (yellow arrows) and *Peak 2* (blue arrows). **(c)** Representative GC-MS EIC for m/z 82.0 of a pheromone-negative control lacking both peaks. *Peaks 1/2* were exclusively detected in pheromone-positive samples. **(b)-(c)** Relative abundance (y-axis) is normalized to the highest signal intensity (100%). The intensity of the highest peak is labeled on each panel’s top-right (highlighted with red triangle). Retention time (x-axis, min) and annotated peaks reflect data acquired with ±0.40 Da mass accuracy in electron ionization positive mode (EI+). Peaks were smoothed using a 2×3 average filter (denoted as Sm (Mn 2×3)). The m/z 82.0 ion corresponds to the target analyte, with the integrated peak area used for quantification. Background signals (e.g., m/z 44.0) are labeled under that peak’s retention times. TOF MS stands for Time-of-Flight Mass Spectrometry. **(d)** Stage- and status-dependent production of *Peaks 1/2* across *C. elegans* and *C. remanei*: pink indicates pheromone production (post-sperm-depletion adults in *C. elegans*; virgin females in *C. remanei*), green indicates lack of pheromone production (larvae, self-sperm non-depleted *C. elegans*, mated *C. remanei* or males), and gray signifies untested conditions.

**Figure 4.**
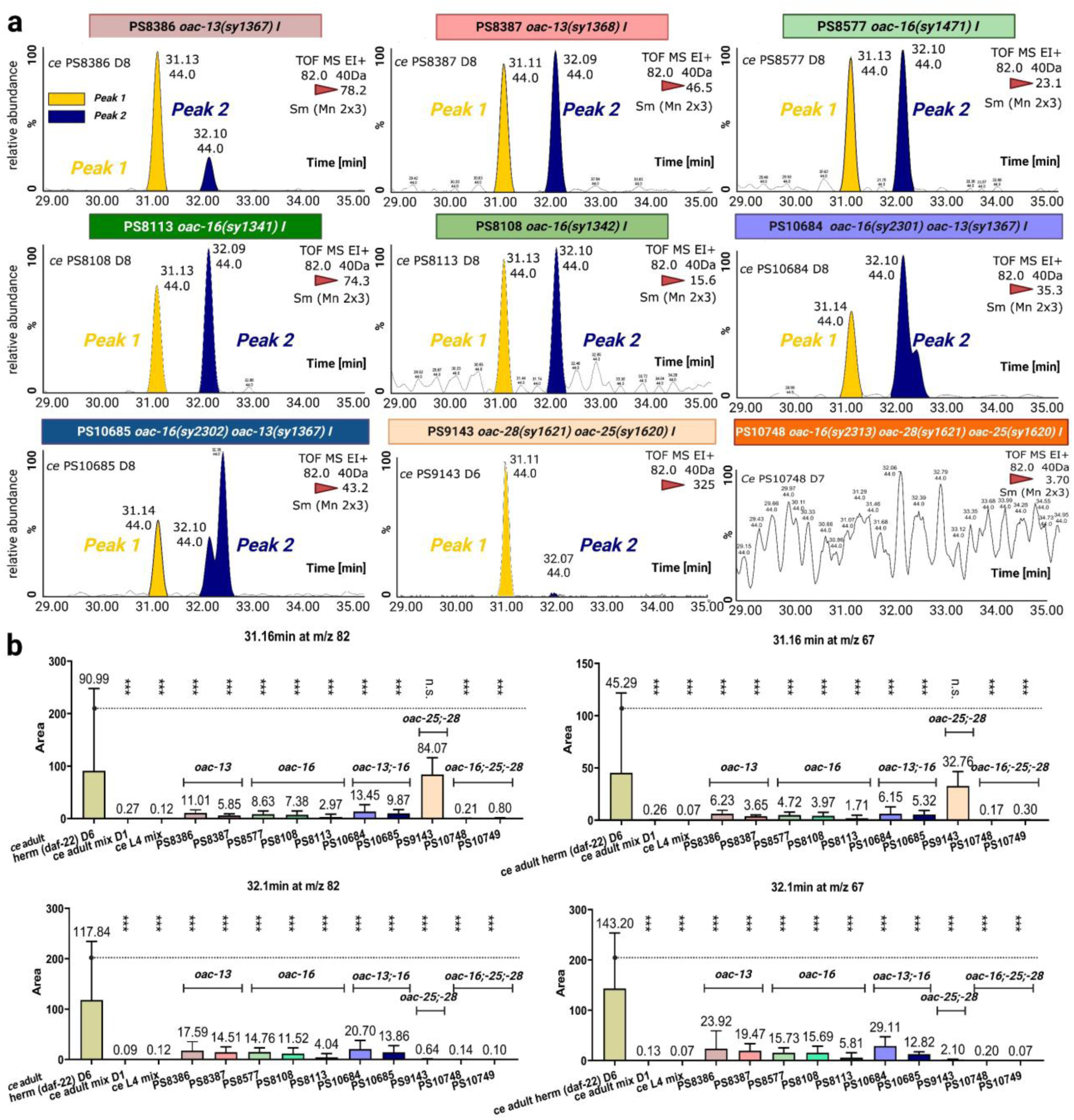
OAC-13, OAC-16, and identical paralogs OAC-25 and OAC-28 are required for the production of volatile pheromone components. **(a)** Representative GC-MS EIC for m/z 82.0 of *(oac-13, oac-16, oac-16 oac-13* double mutant, and *oac-28 oac-25* double mutant). *Peaks 1/2* (t_R_= 31.1 and 32.1 min), identified in wild type pheromone-positive samples (Figure 3), were absent or significantly reduced in all test mutants. *Peaks 1* (yellow arrows) and *Peak 2* (blue arrows)**. (b)** Quantification of integrated peak areas for VSPs-related *Peaks 1/2* (on m/z 82.0 ion and m/z 67.0 ion). Single mutants (*oac-13* and *oac-16*) reduced abundances of both peaks similarly. Double mutants (*oac-16 oac-13)* showed no additive reduction compared with single mutants. *oac-28 oac-25* double mutant abolished *Peak 2* entirely, while *Peak 1* remained unaffected. n = 5–8 biological replicates for each strain. Error bars: S.D.; P < 0.005; P < 0.05 (two-tailed unpaired t-test); ns., not significant (P > 0.05).

Specifically, we extracted pheromones from hermaphrodites lacking self-sperm (to avoid sperm-derived signals that inhibit VSP production; REF) and evaluated their ability to attract wild type males. In particular, we evaluated their activity using a standardized chemoattraction assay with wild type *him-5(e1490)* males. We adopted the pheromone extraction and chemoattraction assay protocol from Wan *et al.* with minor modification ^26^. Of 59 *oac* mutants screened (56 *oac* named genes and *bus-1, nrf-6, rhy-1*) most retained normal pheromone activity (Figure 2a–g). Strikingly, mutants of four genes on chromosome I—two tandem paralogs (*oac-13* and *oac-16*) and two nearly identical paralogs (*oac-25* and *oac-28*)—displayed defective volatile sex pheromone (VSP) production, suggesting impairments in biosynthesis, secretion, or in the availability of a precursor molecule or essential nutrient required for VSP production. *oac-13* and *oac-16* are 179 kilobases apart on the “forward” strand, while *oac-25* and *oac-28* are 6.4 kilobases apart on the reverse strand (Figure 1e). Due to their nearly identical genomic sequences (99% similarity) and close proximity on chromosome I, *oac-25* and *oac-28* are difficult to target individually using standard genetic modification approaches. Furthermore, their coding sequences (CDS) are 100% identical, prompting us to analyze an *oac-28* and *oac-25* double mutant. During the generation of this double mutant, we obtained a single *oac-28* mutant strain and performed pheromone extraction and assays on this strain in parallel.

From the screening, we identified that pheromone extracts from *oac-13* and *oac-16* single mutants and *oac-28 oac-25* double mutants elicited significantly reduced male attraction compared to wild type extracts, whereas the *oac-28* single mutant did not (Figure 3b,c,h). Large-scale trials confirmed these defects (n = 20 trials, 400 males; Figure 3h). Then we evaluated an *oac-16 oac-13* double mutant, which exhibited a similar level of male attraction defect as the cognate single mutants. Since *oac-13* and *oac-16* shows similar male attraction to its double mutant, we tested *oac-16 oac-28 oac-25* triple mutants. Strikingly, this triple mutant almost completely lost male attractiveness (C.I.= 0.19 from two strain, 14 VSP samples and n=800 worms, Figure 2h). Two additional mutants, *oac-49* and *oac-55*, exhibited minor reductions in initial small-scale screening, but these reductions were not statistically significant in large-scale trials (Figure 2h). We then adopted two-choice assays (see Methods) and compared wild type VSP extracts from 6-day-old adult N2 hermaphrodites against those from *oac* mutants. Males (*him-5*) strongly preferred wild type extracts over those from *oac-13* and *oac-16* single mutants, *oac-16 oac-13* and *oac-28 oac-25* double mutants, and *oac-16 oac-28 oac-25* triple mutant (C.I.= 0.705, two strain). These assay results further confirm their pheromone deficiencies (Figure 2i).

OAC-13 and OAC-16 share 60.18% amino acid sequence identity, while OAC-25 and OAC-28 are identical proteins. Further comparisons reveal 70.45% identity between OAC-25/OAC-28 and OAC-16, and 57.78% identity with OAC-13, highlighting significant sequence conservation among these structurally homologous proteins. In addition, all four genes cluster within a conserved genomic region on Chromosome I (13.07–13.20 cM).

### GC-MS analysis identifies two conserved peaks in volatile sex pheromone samples, which are highly regulated by reproductive status

*Caenorhabditis* hermaphrodites produce VSPs only after exhausting their self-sperm, which occurs 5–6 days post-adulthood in wild type individuals or 6–8 days in *oac* mutants. By contrast, virgin adult females of dioecious species continuously secrete these VSPs until mated ^23–27,41^. To identify candidate VSPs, we performed headspace solid-phase microextraction (SPME) coupled with GC-MS (See Methods) on cultures of *C. elegans* and *C. remanei* strains at stages associated with pheromone production (Figure 3, S2).

Pheromone-positive samples have that were tested included (1) 6-day-old adult hermaphrodites of wild type *C. elegans* N2 strain, which were self-sperm depleted; (2) 6-day-old adults of *C. elegans daf-22* mutants, defective in ascaroside biosynthesis, to control for contributions from ascaroside signals in GC-MS; and (3) 1-day-old virgin adult females of wild type *C. remanei*, a dioecious species (Figure 3b, S3). Pheromone-negative controls included (1) 1-day-old adult *C. elegans* hermaphrodites, retaining self-sperm; (2) *C. elegans* larvae (L2–L4 stages), representing non-reproductive developmental stages; (3) *C. remanei* adult males; and (4) mixed-sex 1-day-old adults of *C. remanei*, where mating should suppress the VSP production (Figure 3c, S4). GC-MS analyses revealed two conserved chromatographic peaks (*Peaks 1/2*) present exclusively in all pheromone-positive samples and absent in all negative controls. Despite the unresolved chemical identities of *Peaks 1/2*, the retention time (t_R_) alignment (Figure 3b) and identical mass spectral profiles (Figure 3a, S3a) confirmed their identity across species and strains, suggesting these compounds represent conserved volatile chemicals associated with previously reported male attraction stages (Figure 3d) ^23–27,41^.

### OAC Gene Mutants Exhibit Loss of Volatile Sex Pheromone Signals in *C. elegans*

To determine whether *oac-13, oac-16, oac-25,* and *oac-28* influence VSPs production or secretion, we analyzed pheromone extracts from behaviorally deficient *oac* mutants using SPME-GC-MS. To quantify the results, we analyzed GC elution times, MS profiles, and integrated peak areas derived from extracted ion chromatograms (EICs) for analyte ions (*m/z* 67.0 and 82.0; Figures 4 and S5).

Single mutants of *oac-13* or *oac-16* showed significant reductions in the abundance of *Peak 1/2* compared to wild type controls (Figure 4). The *oac-16 oac-13* double mutant exhibited no further reduction, consistent with behavioral assays, whereas the double mutants impaired male attraction to a similar degree as the single mutant (Figure 2h). By contrast, the *oac-28 oac-25* double mutant retained *Peak 1* but nearly lost *Peak 2* (Figure 4b). While the remaining male attractiveness in *oac-28 oac-25* double mutants suggest that both peaks retain male attractiveness (Figure 2h). The *oac-16 oac-28 oac-25* triple mutant lost both *Peak 1/2*, matching the pheromone-negative controls GC-MS data such as *C. elegans* L4 and one-day-old adults (Figure 4b).

Behavioral assays and biochemical evidence suggest distinct roles of those four OAC genes: *oac-13/oac-16* act upstream, influencing a precursor required for both *Peak 1/2*, while *oac-25/oac-28* function downstream, specifically regulating *Peak 2*. The lack of additive effects in *oac-16 oac-13* double mutants imply they act in a common pathway or complex, with both paralogs essential for synthesizing *Peaks 1/2*. On the other hand, *oac-25/oac-28* have identical protein sequences and are in close chromosomal proximity, which likely share redundant roles in *Peak 2* production.

### *oac-16*, *oac-25* and *oac-28* expression pattern and stage

Transcriptional reporters for *oac-16*, *oac-25*, and *oac-28* were generated to characterize their spatiotemporal expression. All three genes were expressed in hermaphrodite seam cells (specialized lateral line epidermal cells), with *oac-16* and *oac-28* showing constitutive expression from embryogenesis to adulthood. In contrast, *oac-25* expression was restricted to post-embryonic stages, suggesting stage-specific regulation (Fig. 7). The images of the embryo, L1, and L2 stages shown here were captured from the same individual worm throughout its early development (Fig. 7). No differences in expression were detected between hermaphrodites and males, implicating either additional biosynthetic enzymes or sex-specific substrates in VSP production. Technical challenges precluded the generation of a transcriptional reporter for *oac-13*, potentially due to sequence assembly issues in its promoter region sequence. Despite this limitation, the robust seam cell expression of *oac-16*, *oac-25*, and *oac-28* supports a model in which VSPs are synthesized in seam cells.

## Discussion

### Genetic Redundancy and Functional Roles of *oac-13/16* and *oac-25/28* in Volatile Sex Pheromone Production

Our findings demonstrate that four OAC family genes—*oac-13, oac-16, oac-25,* and *oac-28*—are required for the production of volatile sex pheromones. These genes may act by influencing secretion pathways, limiting the availability of essential biosynthetic precursors, or altering the metabolic supply of key nutrients necessary for pheromone generation.

The first model proposes that OAC-13 and OAC-16 mediate production of a shared or distinct precursor utilized in both Peak 1/2 pathways, while OAC-25 and OAC-28 act downstream to specifically convert this precursor into Peak 2. There are two possible explanations for the needs of both OAC-13 and OAC-16. First, the phenotypic similarity between single and double mutants suggests these genes may act sequentially in a shared biochemical pathway. For example, OAC-16 might depend on a substrate produced by OAC-13, or vice versa, therefore, disrupting either pauses the pathway. Alternatively, OAC-13 and OAC-16 may need to form a heteromultimer to enable precursor synthesis. Our second model suggest that OAC-25/OAC-28 might help maintain minimal *Peak 1* production, as losing OAC-13/OAC-16 alone reduces but does not eliminate *Peak 1*, whereas the triple mutant (lacking OAC-16, OAC-25, and OAC-28) loses *Peak 1* entirely. OAC-25 and OAC-28 are nearly identical paralogs, likely arising from a recent tandem duplication event in *C. elegans*. Such sequence identity (100%) and genomic proximity— only 6.4 kb apart on the reverse strand of Chromosome I (Figure 1e)—strongly suggest that these genes are functionally redundant. Genes created by recent duplications often serve as genetic “backups,” providing robustness to essential pathways and buffering against deleterious mutations ^42,43^. In principal they can act as to safeguard an essential process ^44^, consistent with broader principles of genetic robustness ^45–47^.

### Gene duplication in *Caenorhabditis* nematodes

Using BLAST starting with the OAC-25 and OAC-28 protein sequence, we identified homologs across *Caenorhabditis* species and identified hits in *C. remanei*, *C. briggsae*, *C. brenneri and C. nigoni*. No identical proteins were found in closely related species, with the highest sequence matches 54.47% identity to *C. briggsae* proteins: CBG10742. We also used BLAST of OAC-13 and OAC-16 but found no identical proteins and no protein similarity over 60%. While *C. remanei* produces VSPs indistinguishable from those of *C. elegans* by GC-MS retention time and mass spectrometry, we found no highly conserved orthologs of *oac-13, oac-16, oac-25*, or *oac-28* in its genome. This genetic divergence suggests that conserved pheromone components are generated via functionally convergent pathways or a shared catalytic motif.

To characterize the evolutionary duplication event, we performed sequence analysis of 18 orthologous proteins showing >92% query coverage and >50% amino acid similarity to OAC-25/OAC-28. As demonstrated in Figure S6, these proteins display conserved duplication patterns across multiple species. In *C. remanei*, we identified two highly conserved paralog pairs (96.2% and 98% identical). *C. briggsae* showed even greater conservation, with one paralog pair exhibiting 99.6% similarity and three additional paralogs demonstrating near-perfect identity (100% and 99.6%). All mapped genes are located near their paralogs in the genome. There are three genes that have not been mapped yet. These levels of sequence conservation strongly suggest recent, lineage-specific gene duplication events in both species. In our analysis, only the double mutant of *oac-25* and *oac-28* affects the VSPs synthesis, consistent with the dual role of gene duplication in evolution: maintaining stability while enabling innovation ^43,48,49^. Future investigations in nematodes across the evolutionary tree-derived species (both ancestral to and descendant from the *C. elegans* lineage) could clarify whether OAC-25 and OAC-28 function as redundant backups or sub-functionalized intermediates, which would elucidate their evolutionary trajectory and distinguish between conserved redundancy (shared ancestral function) and lineage-specific specialization ^50^.

### Volatile sex pheromone biosynthesis location

Prior work established that laser ablation of Z1 and Z4—precursor cells of the somatic gonad ^51^—abolishes hermaphrodite male attractiveness, while ablation of germline precursors Z2 and Z3 has no effect, implicating the somatic gonad in volatile pheromone (VSP) biosynthesis ^24^. Consistent with this finding, vulvaless hermaphrodites retain wild-type attractiveness, ruling out the vulva as a secretion opening for VSPs ^25^. Our expression analysis of *oac-16*, *oac-25*, and *oac-28* localizes these genes to seam cells (Fig. 5). We thus infer that the seam cells are a crucial site of VSP biosynthesis and might be the source of secreted VSP.

**Figure 5.**
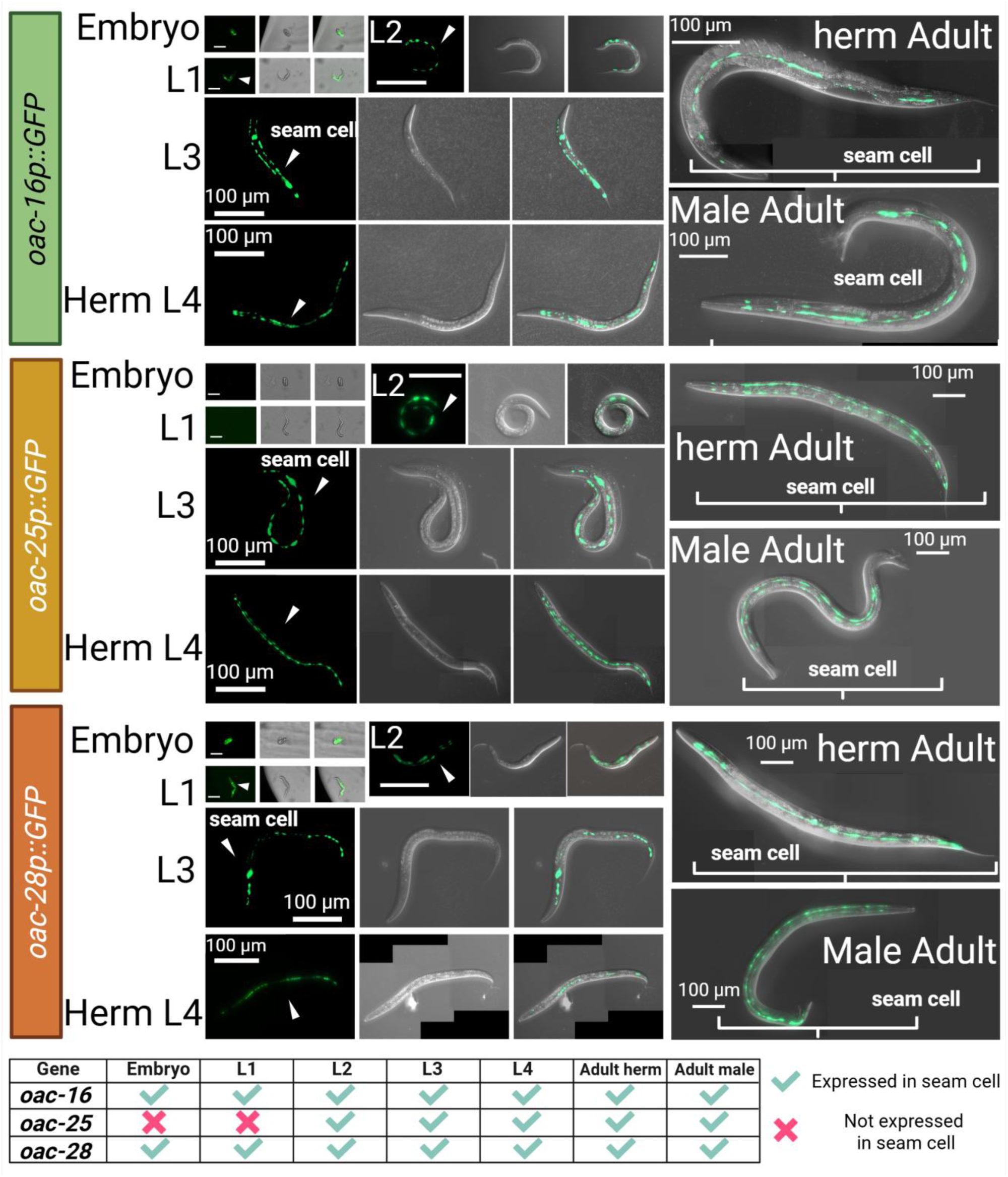
Volatile sex pheromone related OAC stage specific expression pattern. We constructed transcriptional reporters by cloning the *oac-16* (PS10804)*, oac-25* (PS10805), and *oac-28* (PS10806) promoters upstream of gfp in the pPD95.75 vector (see Methods). All three OAC genes are expressed in seam cells. The embryo, L1, and L2 images shown here are from the same individual worm. Stable extrachromosomal lines were established.

The somatic gonad of *C. elegans* hermaphrodites comprises eight distinct structural regions, including connections to the vulva and seam cells of the lateral epidermis, the uterus, spermathecal valve, spermatheca, oviduct, ovary sheath, and distal tip ^52,53^. Notably, the uterine-seam (utse) syncytium forms a critical physical linkage between the somatic gonad and hypodermal seam cells during the mid-to-late L4 stage ^52,54^. These anatomical and developmental insights contextualize our findings on VSPs biosynthesis. The L4-stage establishment of utse cell connectivity coincides with the onset of VSP production ^24^, raising the possibility that somatic gonadal signaling to seam cells—mediated through this physical linkage—may regulate pheromone synthesis. While VSPs are not produced in the gonad itself, the gonad’s structural integration with seam cells (via the utse cell) may provide developmental cues to modulate OAC gene activity in seam cells. For instance, the gonad could relay other metabolic or temporal signals during late larval stages to initiate or terminate VSPs production. Future work could assess whether disrupting utse-seam cell connectivity (e.g., via mutations in adhesion genes) ablates VSPs synthesis, linking morphogenesis to pheromone regulation.

The lateral position and established secretory functions of the seam cells make them exceptionally well-suited for synthesizing and secreting volatile pheromones. Embedded within the hypodermis but exposed to the external environment via their apical surface directly underlying the cuticle (including the alae) ^55–57^, seam cells possess the physical location necessary for efficient volatile compound release. Furthermore, their documented capacity for synthesizing and secreting stage-specific cuticular components, such as collagens and the specialized alae structures ^58^, demonstrates robust biosynthetic and secretory machinery. This existing molecular infrastructure for producing extracellular molecules aligns with a role in VSP biosynthesis and direct secretion into the surrounding environment.

### Conservation of volatile sex pheromones across nematode lineages with divergent reproductive strategies

*Peaks 1/2* were detected in pheromone-positive samples of both *C. elegans* (androdiecious, hermaphrodites/males) and *C. remanei* (dioecious, females/males), despite their phylogenetic divergence (∼100 million years) and divergent sexual systems. The presence of *Peaks 1/2* in both suggests these VSP signals evolved early before their sexual systems diverged. A key question is how species avoid cross-attraction. Spatial separation of populations, differences in pheromone concentration and components ratio, or other species-specific ascarosides may help maintain reproductive isolation ^59–62^. Further comparing *Peaks 1/2* across nematode species could reveal whether their ratios or combinations drive speciation.

### Evolutionary Implications of gene family expansion

Complete loss-of-function alleles are the gold standard for genetic studies. O-acyltransferases are used in many different pathways throughout the cells of eukaryotic animals, and we now have the tools to elucidate their functions in *C. elegans*. Our creation of null mutants for the entire OAC gene family allows for future studies of individual genes or the family as a whole. It is particularly interesting to study why this gene family is largely expanded in *Caenorhabditis* and how they contribute to species-specific pheromone signaling and lead to species differentiation.

All identified *oac* genes associated with VSPs synthesis are clustered closely on chromosome I and exhibit close phylogenetic relationships. *oac-4 is* evolutionary related to those four VSPs-synthesis-associated OAC proteins with high sequence similarity (Figure 1a). In contrast, *oac-4* is the sole gene not required for VSPs synthesis and is located on a distinct chromosome. Sequence analysis of the *oac-4* shows a unique insertion that contains two repeat HWPIYA[F/H] fragment insertions in the SYH catalytic motif (highlighted with pink boxes in Figure S1), a feature absents in all other 58 *C. elegans* OAC family proteins. These insertions may disrupt *oac-4*’ s ability to synthesize VSPs, suggesting potential repurpose of this gene for novel functional roles.

The *Caenorhabditis* genus’s OAC expansion contrasts with other nematodes, suggesting lineage-specific adaptations. The conserved SYH catalytic motif across OACs indicates a shared enzymatic mechanism, while domain architectures (e.g., SGNH hydrolase or NRF domains) suggest divergent functions. For instance, *oac-13* and *oac-16* both essential for *Peak 1/2* may reflect neofunctionalization after duplication, and two identical genes *oac-25* and *oac-28* ensure the *peak 1* production may reflect gene function redundance. All four OAC genes involved in the same VSPs synthesis pathway are evolutionally closed evidenced by phylogenetic clustering and sequence alignment, also their genomic loci are remarkably close.

## Materials and Methods

### Protein alignment

Protein alignments were performed using the ClustalOmega multiple sequence alignment tool with default settings. Protein sequences were obtained from WormBase (WS295; https://wormbase.org).

#### Phylogenetic analysis of OAC proteins

Protein sequences of *Caenorhabditis elegans* OAC family members were retrieved from WormBase WS295. The multiple sequence alignment was performed using the MUSCLE algorithm in MEGA (Molecular Evolutionary Genetics Analysis, version 12) with default parameters. The alignment was trimmed to exclude positions containing gaps or missing data, retaining conserved domains to ensure phylogenetic robustness. A maximum likelihood (ML) phylogeny was inferred using the Tamura-Nei model ^63^. Tree topology was refined using the Nearest-Neighbor-Interchange (NNI) heuristic method, with initial trees generated automatically via default neighbor-joining (NJ) and maximum parsimony (MP) algorithms. Branch lengths represent substitutions per site, and the final tree was visualized and annotated in MEGA.

### *C. elegans* strain construction and maintenance

The Bristol N2 strain was used as a wildtype for *C. elegans*. The *rhy-1*, *bus-1*, and *nrf-6* strains were obtained from the *Caenorhabditis* Genetics Center (CGC). Supplemental Table 3 shows a list of all strains used in this paper. All nematode strains were kept on NGM agar plates seeded with *E. coli* (OP50) at 20°C.

The creation of deletion mutants by non-homologous end joining was done using a modification of the method described in Köhler *et al.* (2017) ^64^ in which only one guide RNA was used. Supplemental Table 1 gives the flanking sequences around the deletions and the primers used in the creation of strains using the single-primer deletion method.

Creation of triple-stop knock-in mutants was done using the universal STOP-IN cassette method as described in Wang *et al.* (2017) ^65^. Supplemental Table 2 gives the flanking sequences around the insertions and the primers used in the creation of strains using the triple-stop knock-in method.

#### Volatile Sex Pheromone extraction

VSPs were extracted from synchronized *C. elegans* hermaphrodites and *C. remanei* females as described in Wan *et al.* 2024 with minor modifications ^26^. For the chemoattraction assay, use 100 µL of solution containing 100 *C. elegans*. For each GC–MS analysis, 2 mL of VSP extract was used per sample. Extracts were prepared from 100 µL of *C. elegans* worm pellet soaked in 2 mL of M9 buffer. For *C. remanei* GC–MS analyses, samples were prepared from 120 individuals soaked in 2 mL of solution.

Worms were synchronized via bleach lysis, washed in M9 buffer. The extraction day was determined by the exhaustion of self-sperm, with no more new-born progeny presenting on the NGM plate. On the extraction day, worms were incubated for 6 hours in M9 to allow pheromone accumulation. Following centrifugation (15,000 × g, 30–60 s), the supernatant containing pheromones was stored at −80°C.

#### Chemoattraction assay and two-choice assay

Male chemotaxis responses were quantified using a chemoattraction assay as described in Wan *et al.* 2024 ^26^. Briefly, 2 µL of pheromone extract or M9 control was applied to designated spots alongside 1 M sodium aside to immobilize arriving males. 20 one-day-old adult males were released at the starting point, and chemotaxis indices (C.I.) were calculated after 30 minutes using: C.I.= [(E−C)/(E+C+N)]. E, C, and N represent worms at experimental, control, or neither spot, respectively. The two-choice assay was performed using the same setup as the chemoattraction assay. The M9 control buffer was replaced with wild type N2 *C. elegans* sex pheromone extract. Chemotaxis indices (C.I.) were calculated using C.I.= [(WT_VSP – OAC_VSP Mutant)/(WT_VSP + OAC_VSP Mutant +N)]. WT_VSP, OAC_VSP Mutant, and N represent worms at wildtype pheromone, oac mutant pheromone, or neither spot, respectively.

#### Solid-Phase MicroExtraction (SPME)

Volatile compounds were collected from hermaphrodite cultures using headspace solid-phase microextraction (SPME). A divinylbenzene/carboxen/polydimethylsiloxane (DVB/CAR/PDMS, 50/30 µm) fiber (Supelco, Bellefonte, PA) was preconditioned for 10 min at 265°C in the GC injector before each use. For sampling, the fiber was exposed to the headspace of sealed culture vials for 10 min at 60°C under continuous agitation (250 rpm, 5 s-on /20 s-off cycles) using a CTC Analytics autosampler. After extraction, fibers were immediately desorbed in the GC injector for 6 min.

#### Gas Chromatography-Mass Spectrometry (GC-MS)

Analyses were performed on an Agilent 6890N gas chromatograph coupled to a Waters GCT Premier orthogonal acceleration time-of-flight mass spectrometer (oaTOF-MS). Separation was achieved using a DB-624 capillary column (30 m × 250 µm × 1.40 µm; Agilent). Helium carrier gas was maintained at a constant flow rate of 0.80 mL/min. The oven temperature program was initiated at 25°C (0 min hold), ramped at 5°C/min to 85°C (0 min hold), then ramped at 1°C/min to 150°C (0 min hold), followed by a 30°C/min ramp to 250°C (1.5 min hold). Electron ionization positive (EI+) was performed at 70 eV, with mass spectra acquired in positive ion mode over a range of m/z 35–500. Blank runs (SPME fiber exposed to miliQ H_2_O sample vials) were interspersed with samples to monitor carryover. System performance was validated daily using alkane standard mixtures (C7–C30). All biological replicates (n = 5–8 per condition) were analyzed in randomized order to minimize batch effects.

#### Data acquisition and quantification

Raw data were processed using Waters MassLynx software. *Peaks 1/2* (putative pheromone components) were quantified by integrating ion chromatograms (m/z 67.0 ± 0.3 and m/z 82.1 ± 0.3) within a retention time window of ±0.4 min around predicted values (t_R_=31.1 min and 32.1 min, respectively). External calibration curves were generated using absolute peak areas, with baseline correction and noise thresholds applied uniformly across samples Peaks 1 and 2 were identified across samples based on retention time alignment (+-0.1% tolerance) and spectral matching.

### Expression pattern imaging

Transcriptional reporters were constructed in vector pPD95.75, driving GFP expression from the *oac-16p*, *oac-25p* or *oac-28p* promoters (PS10804: *him-5;* syEx2004[pPD95.75::*oac-16p*::gfp], PS10805: *him-5;* syEx2005[pPD95.75::*oac-25p*::gfp], PS10806: *him-5;* syEx2006[pPD95.75::*oac-28p*::gfp]). Plasmid DNA (80 ng/µL) was co-injected with the coelomocyte marker *unc-122p*::RFP (20 ng/µL) into *him-5(e1490)* mutants to obtain males for imaging. Transgenic lines expressing these extrachromosomal arrays were isolated, maintained and imaged.

We cultured *C. elegans* in a 398-well plate format, with one embryo per well, to enable imaging at the embryo, L1, and L2 stages. Worms were immobilized using Levamisole, imaged within the wells, and then washed out, which prevented recovery of the same individuals for subsequent time points. Consequently, worms imaged at later stages (L3 to adult) were cultured separately using a different method optimized for those stages. Although this limits continuous tracking of the same individual throughout development, the use of stage-matched populations and consistent imaging conditions allows for reliable comparisons across stages. Consequently, worms imaged at later stages (L3 to adult) were cultured separately using a different method optimized for those stages. For these stages, animals were immobilized on a 4% ultrapure agarose pad in 5 mM Levamisole (in H₂O) and imaged using a Zeiss AxioImager2 microscope equipped with a Colibri 7 LED fluorescence illumination system and an Axiocam 506 Mono camera (Carl Zeiss Inc.).

## Author contributions

X.W. conceptualized and performed investigation (sample extraction, behavioral assays, GC-MS experiments); S.M.C. and H.P. designed and generated *oac* mutant strains with help from M.T.; Y.Y. and S. M. C. conducted metabolism experiments and analysis; A. G. designed and generated the *oac* reporter strains and R.M. performed the imaging. J. L and X. W. performed phylogenetic and sequence analysis. G.R.M., S.R. and X. W. performed the initial pheromone GC-MS study. X.W. and P.W.S. wrote the original draft; All authors reviewed and edited the manuscript; F.C.S. and P.W.S. supervised the project.

## Acknowledgments

We acknowledge Nathan F. Dalleska and the Caltech Resnick Water and Environment Lab for the GC-MS analysis, and King L. Chow and the HKUST Environmental Central Facility for the initial GC-MS study. We thanks Jessica J. Sun for the initial phylogenetic tree analysis. We thank Joshua N. Muller for assisting with the imaging process. We also thank Erich Schwarz and Jae Cho for their assistance with WormBase data mining. We thank the *Caenorhabditis* Genetics Center (CGC), which is funded by the National Institutes of Health (NIH), for some *C. elegans* strains used in these experiments. This work was supported by the National Science Foundation Graduate Research Fellowship under Grant No. DGE 1745301 to S.M.C., a Caltech SURF to J.J.S., and the NIH under Grant R24OD023041 to P.W.S, a Bren Professor of Biology. Tianqiao and Chrissy Chen Institute for Neuroscience senior postdoc fellowship and Tianqiao and Chrissy Chen Institute for Neuroscience postdoc innovator grant to X.W. We thank the Chuck Lorre Research Scholars Program for supporting R.M.

## Supplemental Information

**Supplemental Figure 1.**
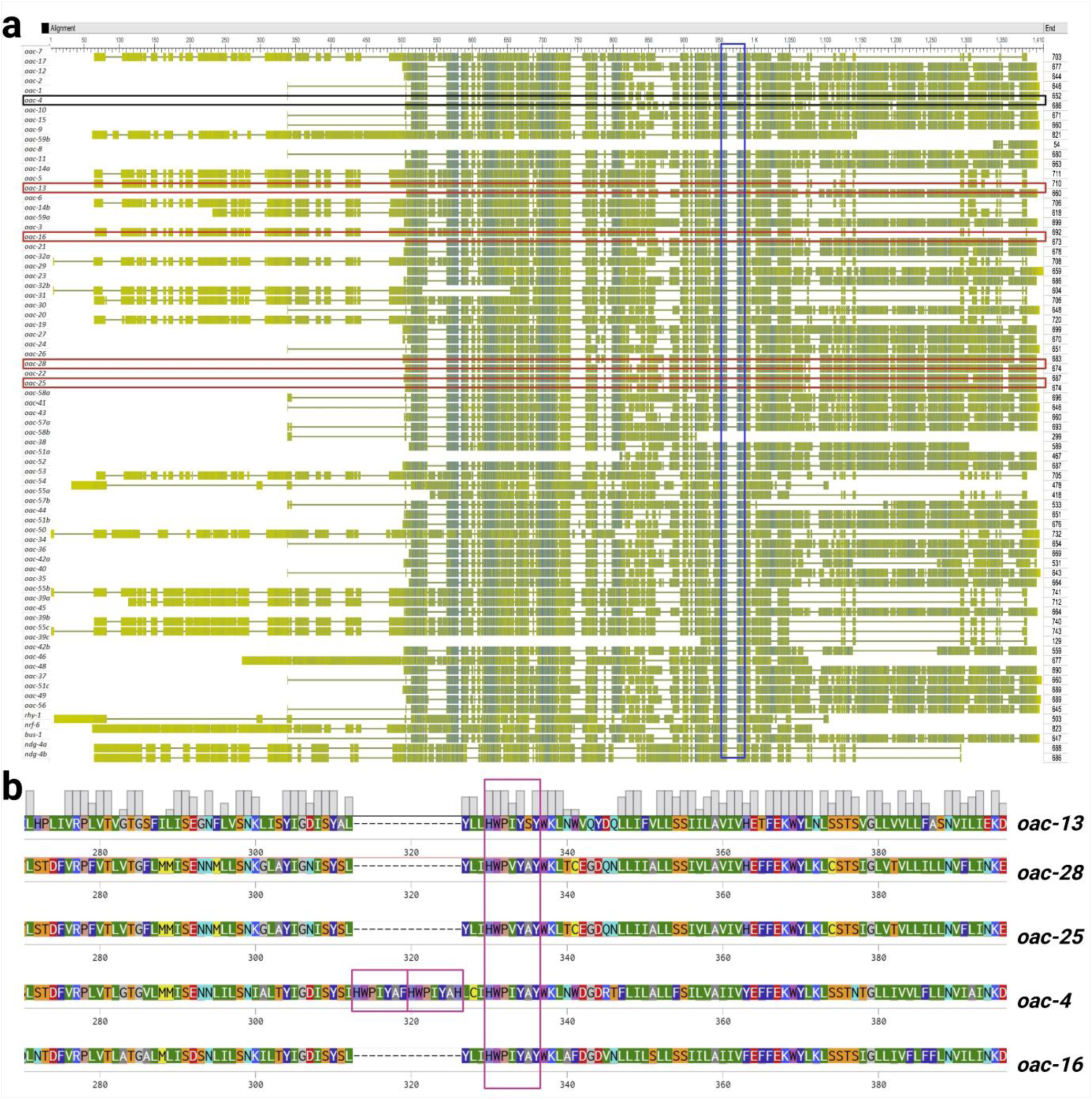
Conserved SYH catalytic motif in OAC acyltransferase 3 domains. **(a)** Sequence alignment of OAC proteins with acyltransferase 3 domains, visualized using BLOSUM substitution matrices. Residue conservation is color-coded: blue = high similarity, green = low similarity. Red boxes highlight *oac* family genes linked to VSPs synthesis in this study; black box denotes *oac-4*. The SYH catalytic motif (blue box) is conserved across all OAC family members. **(b)** The protein sequence of the *oac-4* insertion sequence within the SYH catalytic motif. *oac-4* uniquely contains two HWPIYA[F/H] fragment insertions (pink boxes) within this motif. Multiple sequence alignment of OAC proteins involved in VSP synthesis and the evolutionary related gene *oac-4*.

**Supplemental Figure 2.**
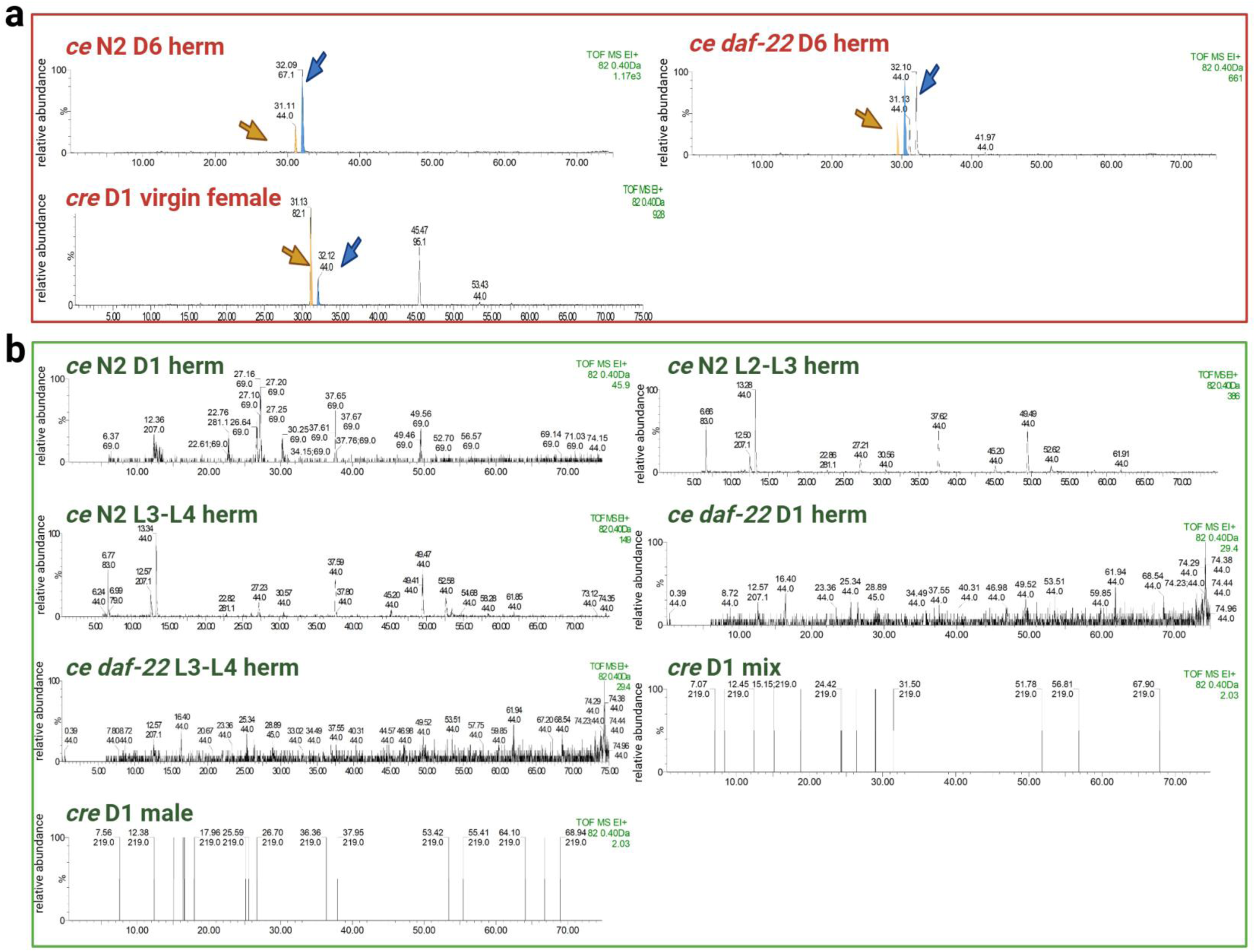
Full GC-MS EIC for *m/z* 82.0 of pheromone-positive and pheromone-negative samples. *Peaks 1/2* were exclusively observed in pheromone-positive samples. **(a)** Representative EIC (0– 75 min) of pheromone-positive samples, including 6-day-old self-sperm-depleted *C. elegans* N2 (wild-type) and *daf-22* mutant hermaphrodites, and 1-day-old unmated *C. remanei* females. *Peaks 1* (yellow arrows, t_R_=31.1min) and *peak 2* (blue arrows, t_R_=32.1min) are characteristic of pheromone presence. **(b)** Representative GC profile (0–75 min) of a pheromone-negative control lacking both peaks.

**Supplemental Figure 3.**
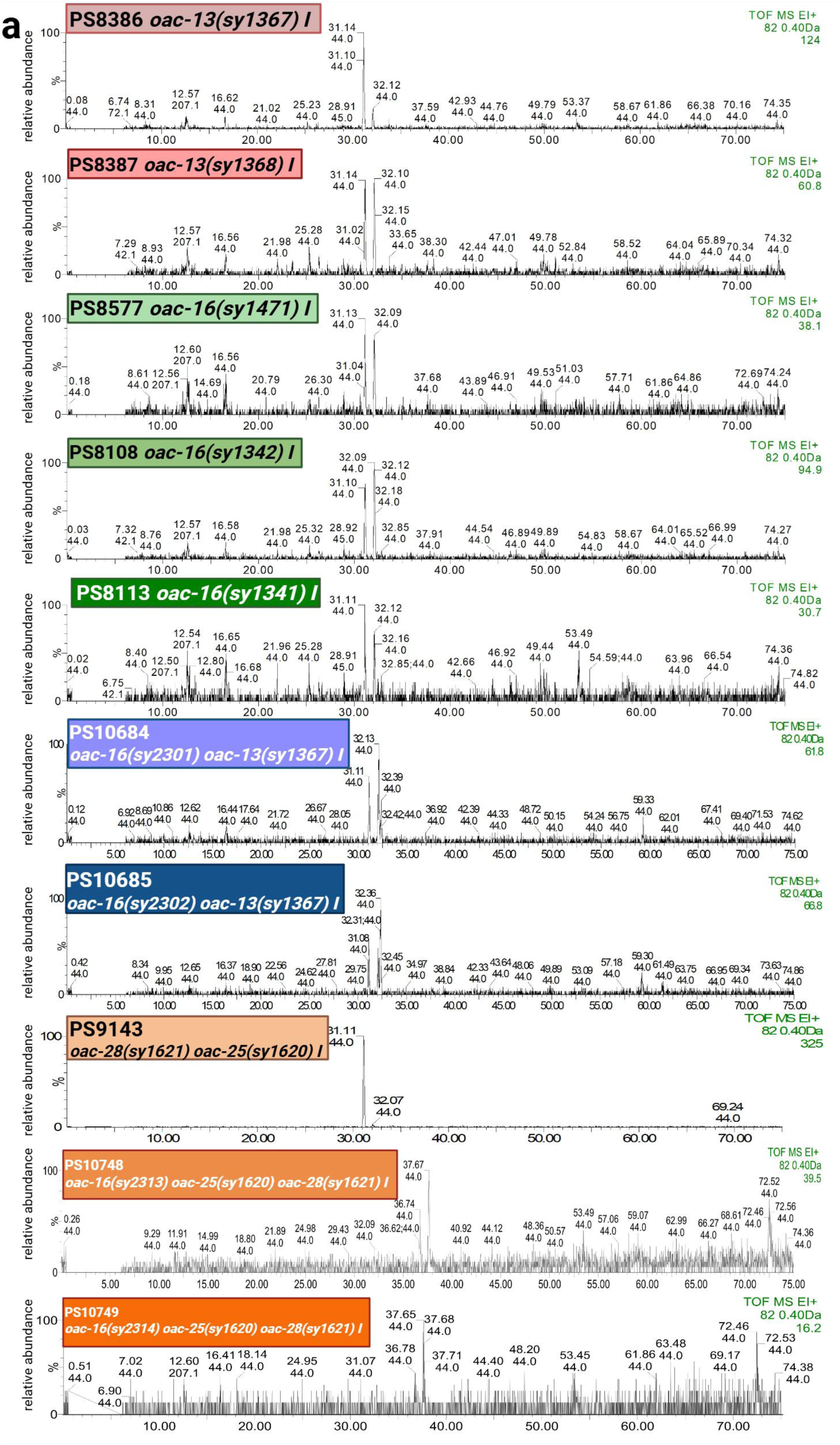
**(a)** Full MS (0-500 m/z) of *Peak 1* (t_R_=31.1 min) and *Peak 2* (t_R_=32.1 min) detected in pheromone-positive samples. Retention times and mass spectra were conserved across samples. **(b)** GC profiles (0–75 min) of VSPs positive data: 6-day-old N2 hermaphrodites, showing EICs for m/z 67.0 and m/z 82.0, alongside the total ion chromatogram (TIC).

**Supplemental Figure 4.**
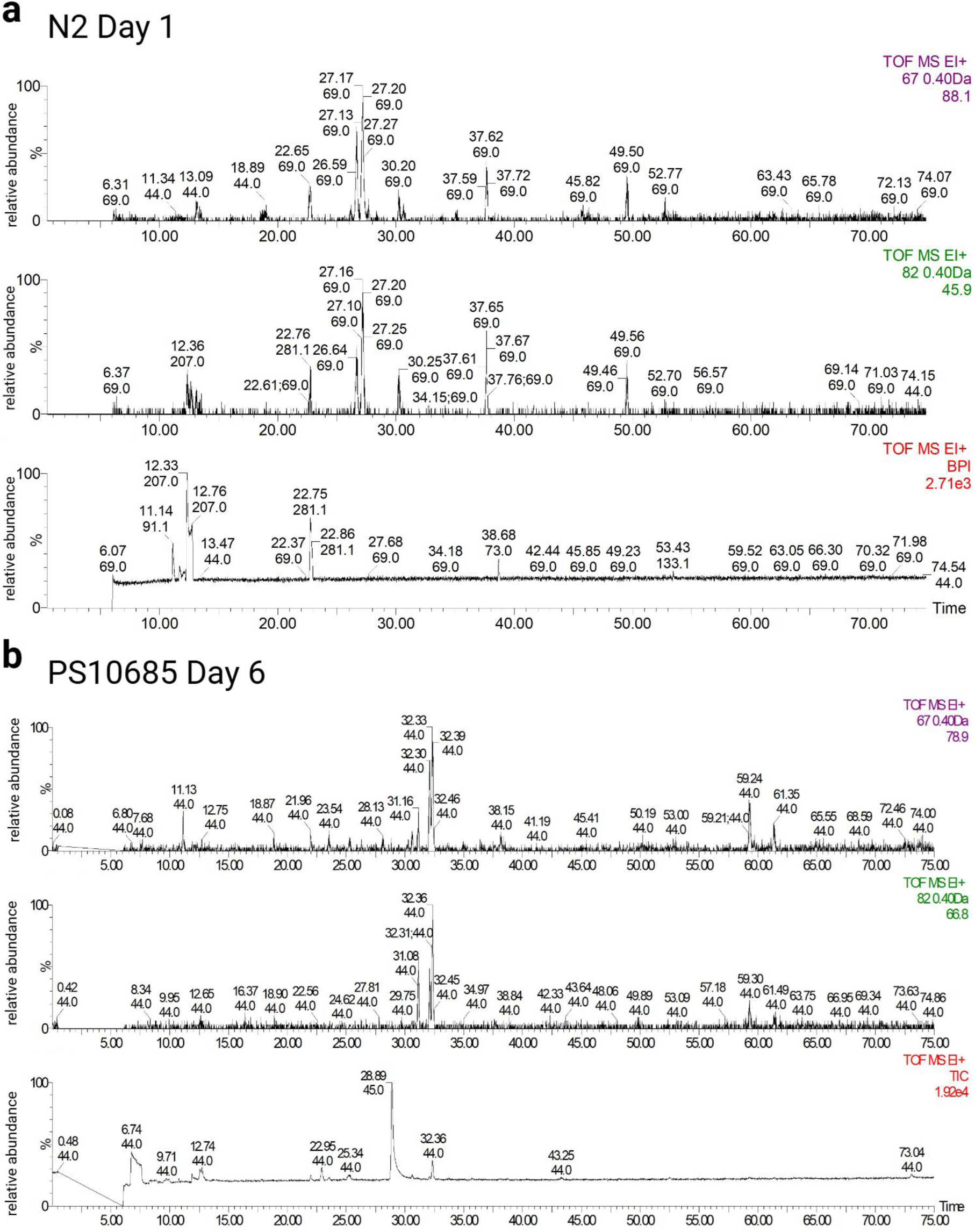
GC profiles (0–75 min) of VSPs negative data: 1-day-old N2 hermaphrodites and 6-day-old *oac-16 oac-13* double mutant hermaphrodites showing EICs for m/z 67.0 and m/z 82.0, alongside TIC.

**Supplemental Figure 5.**
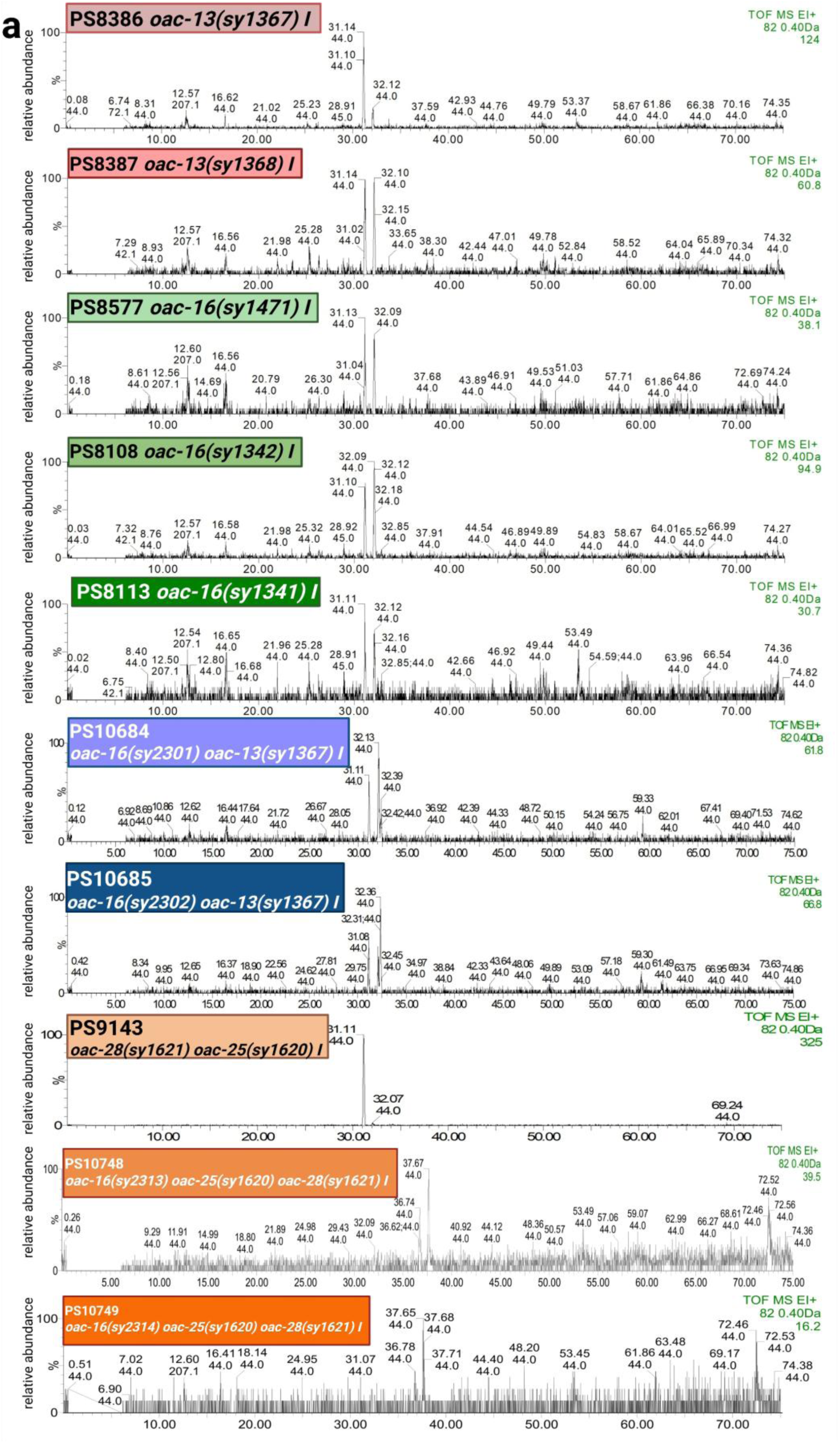
Full GC profiles (0–75 min; m/z 82.0 ion) of *oac-13, oac-16, oac-16 oac-13* double mutant, and *oac-28 oac-25* double mutant. *Peaks 1/2* (t_R_=31.1 min and 32.1 min), corresponding to pheromone components identified in wild type samples (See Figure 3 and S2), were abolished or markedly reduced in all mutant strains. *oac-13, oac-16* single mutant, and *oac-16 oac-13* double mutant reduced abundances of both peaks similarly. *oac-28 oac-25* double mutant abolished *Peak 2* entirely, while *Peak 1* remained unaffected. (n = 5–8 biological replicates)

**Supplemental Figure 6.**
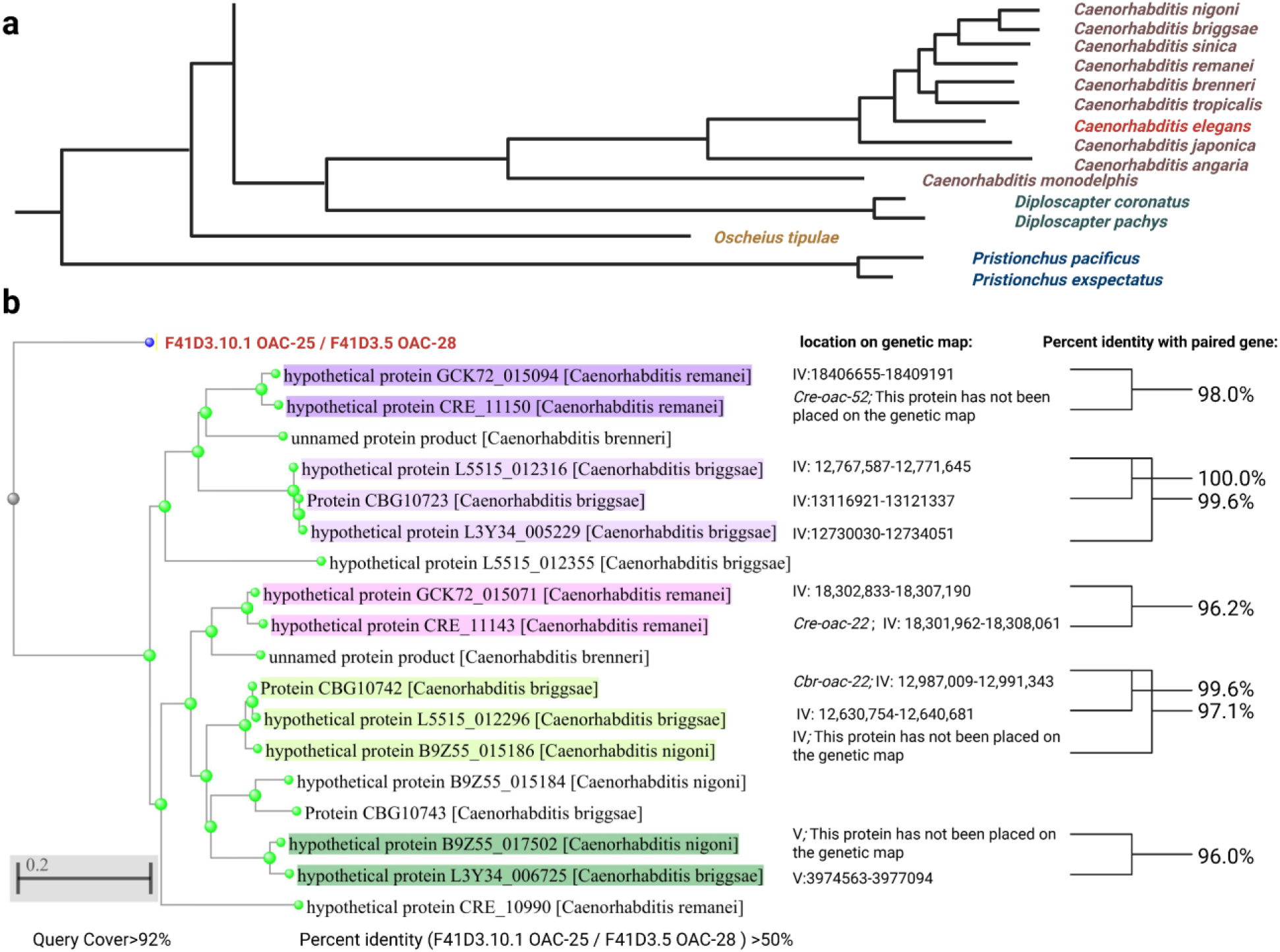
Lineage-specific gene duplication of OAC-25/OAC-28 homologs occurred within the *Caenorhabditis* lineage prior to and following the divergence of *C. elegans*. **(a)** Phylogenomic map of *Caenorhabditis* Species, *Diploscapter* Species*, Pristionchus* Species, and *Oscheius tipulae*. Figure modified from ^66–68^ **(b)** BLAST analysis of OAC-25/OAC-28 protein sequences in nematode taxa diverging prior to, and subsequent to, the emergence of *C. elegans* shows no identical proteins were detected, with maximum sequence identities of 55.39% (*C. remanei* hypothetical protein GCK72_015070) and 54% (*C. briggsae* CBG10742 and L5515_012296). Among 16 proteins with >92% query coverage and >50% identity to OAC-25/OAC-28, there are two paralog pairs in *C. remanei* and one pair plus three near-identical paralogs in *C. briggsae.* Paralogous protein pairs are highlighted with corresponding colors, while genomic locations and percentage identity values relative to their paralogous counterparts are annotated in the right panel. This analysis and accompanying figure were produced using the NCBI BLASTP tool.

**Supplemental Table 1.**
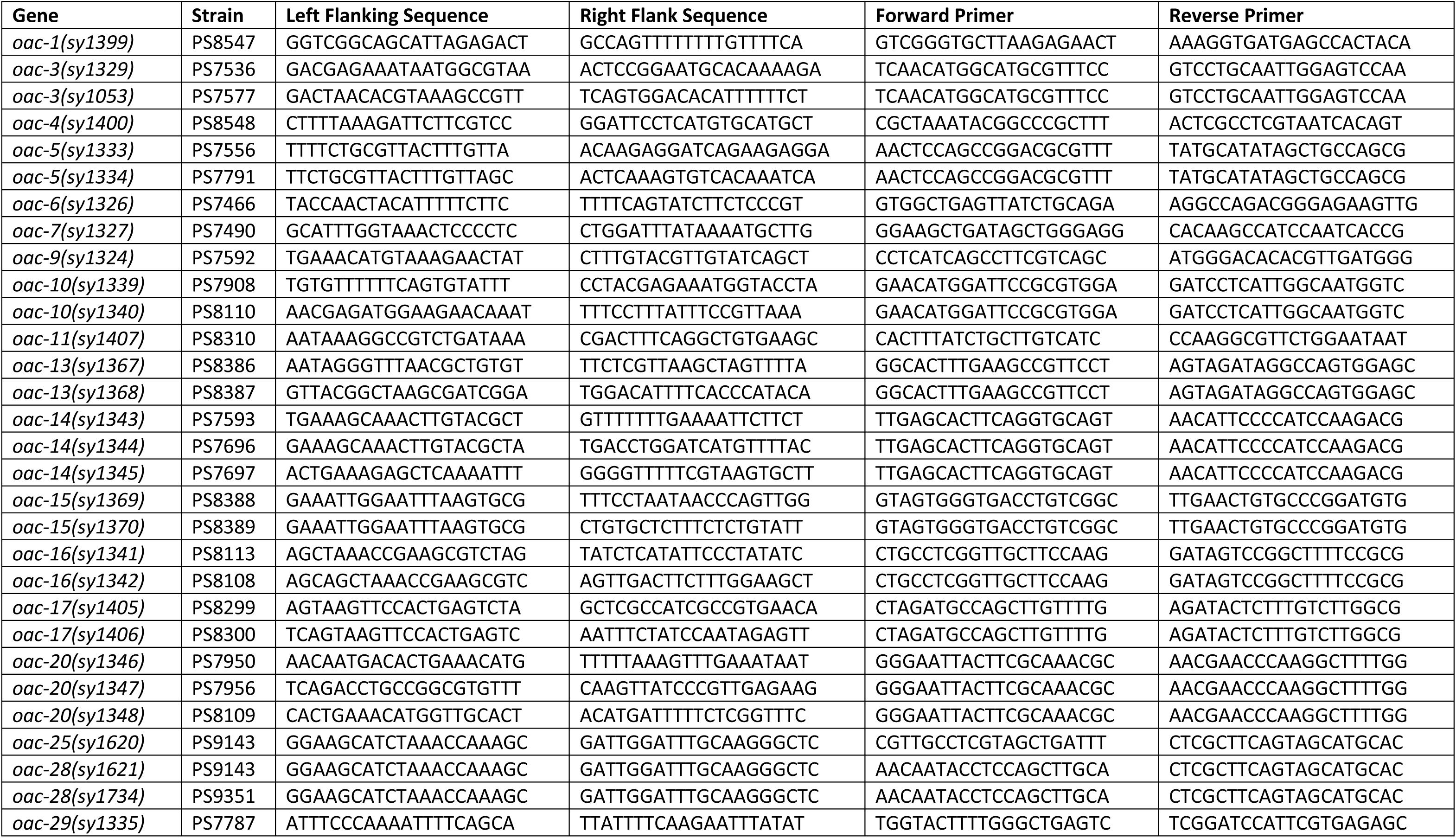
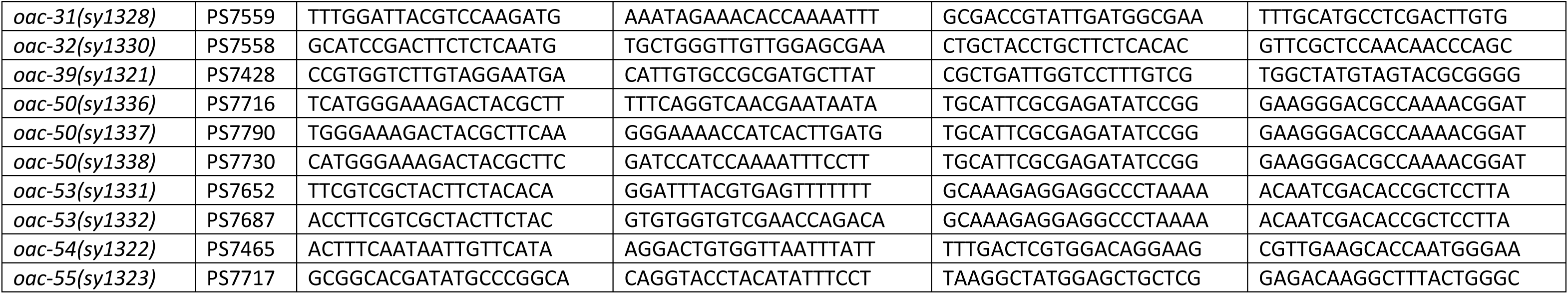
List of all *oac* mutant strains made by deletion method.

**Supplemental Table 2.**
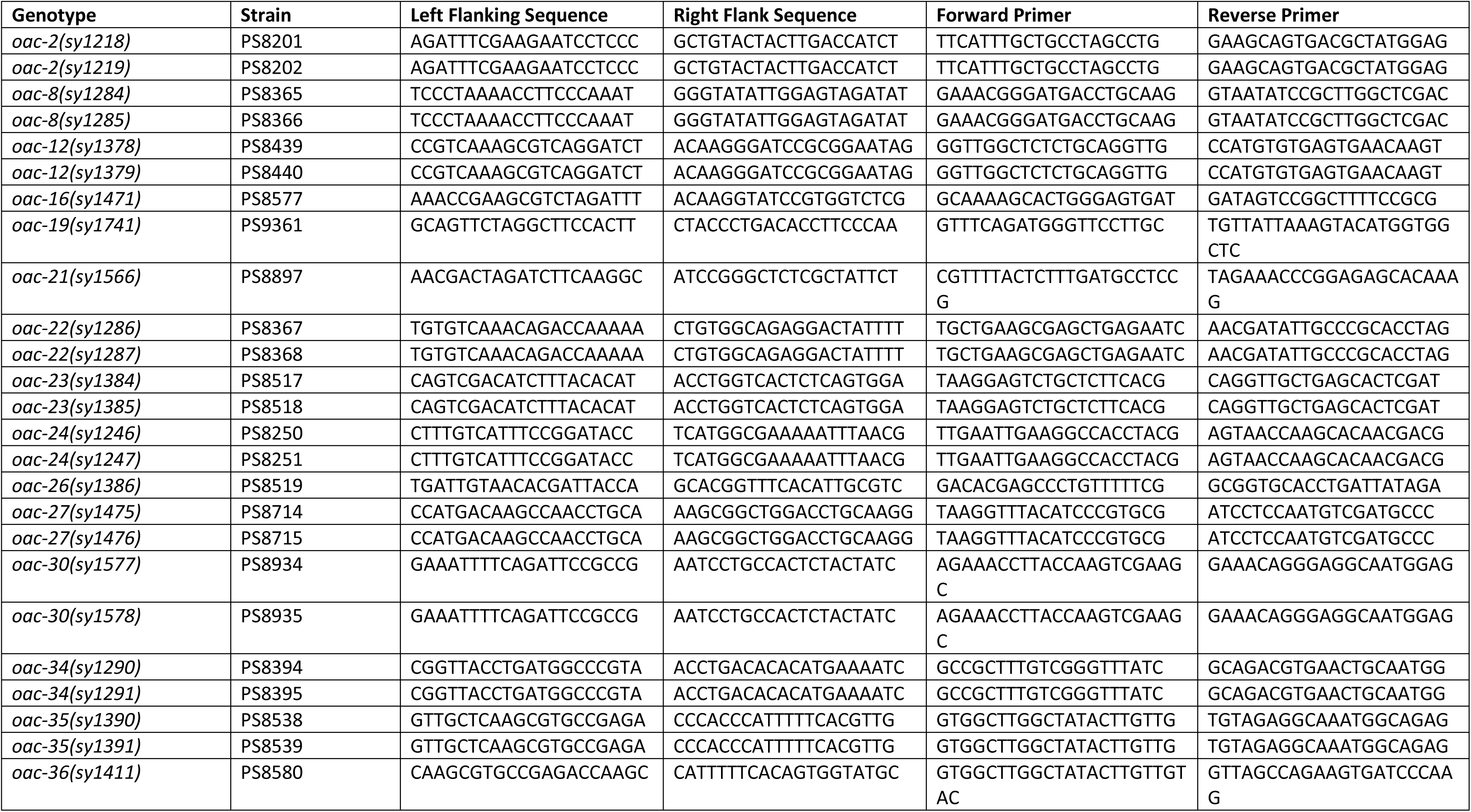
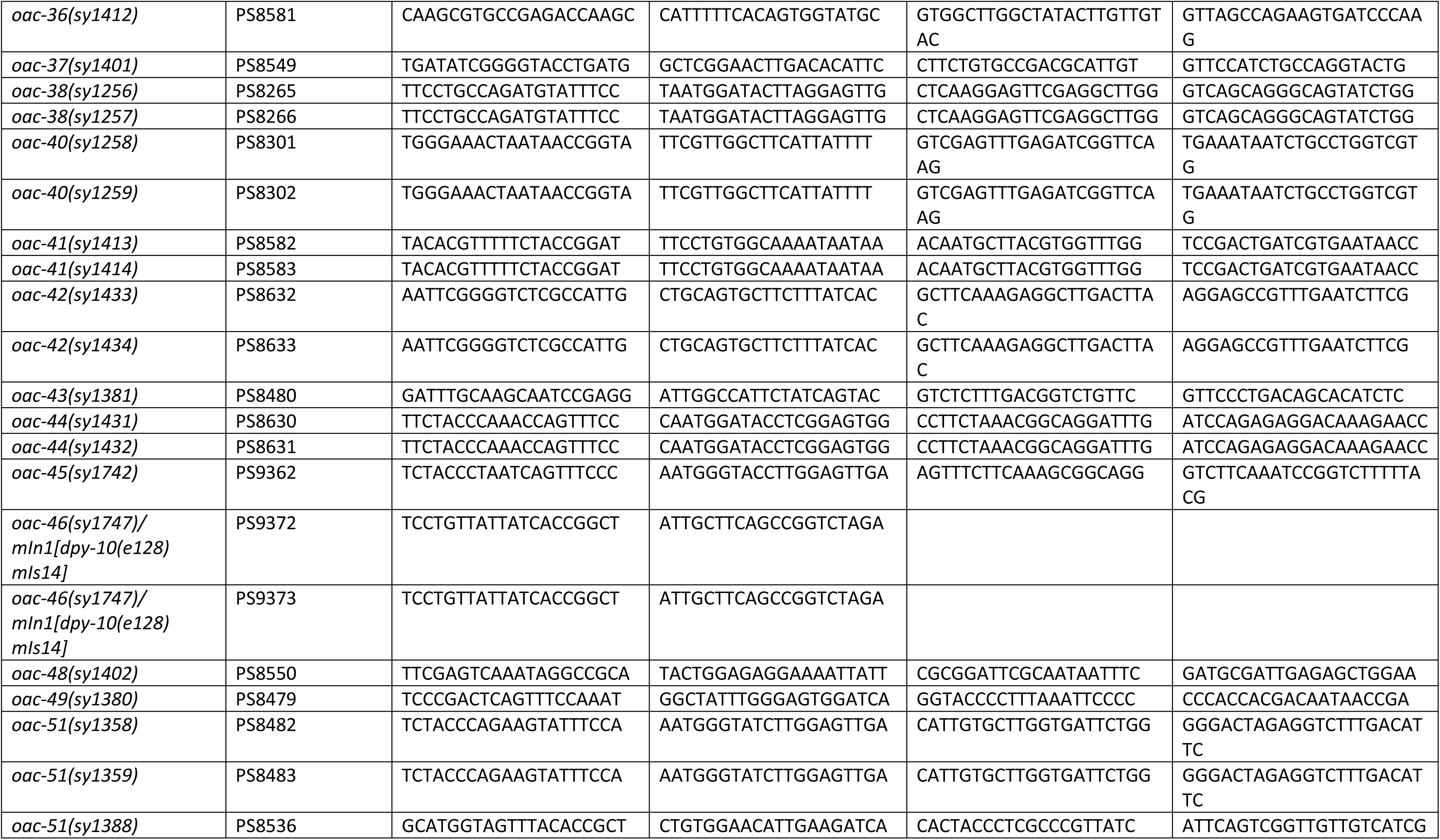
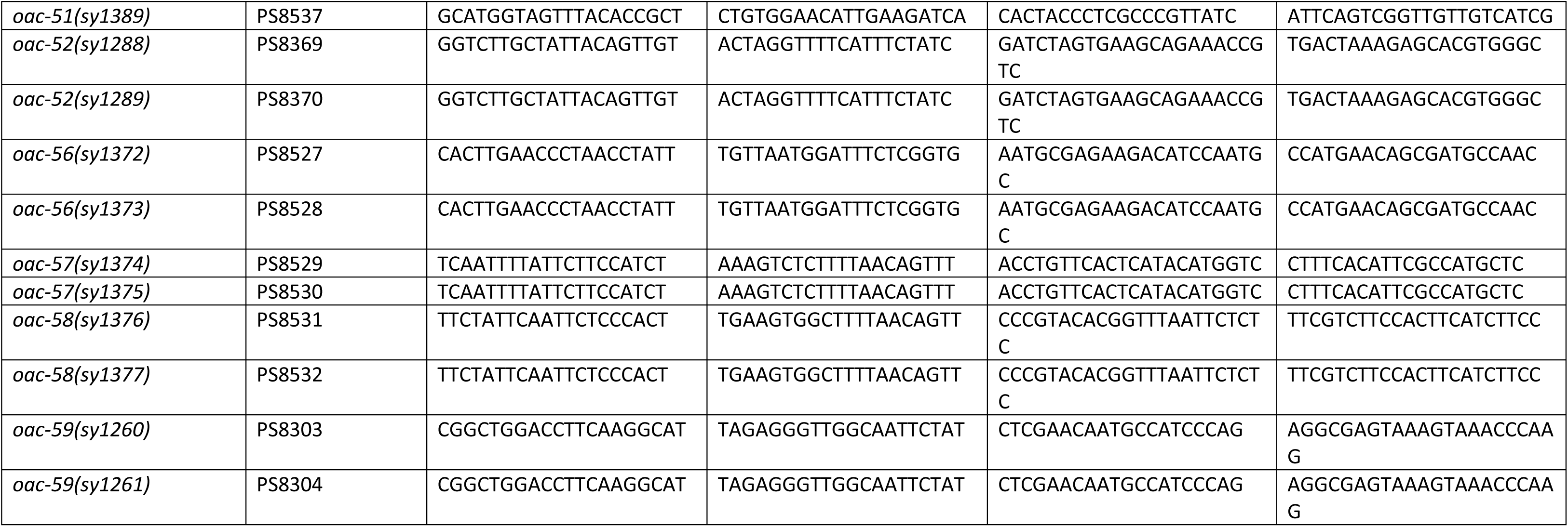
List of all *oac* mutant strains made by insertion method.

**Supplemental Table 3.**
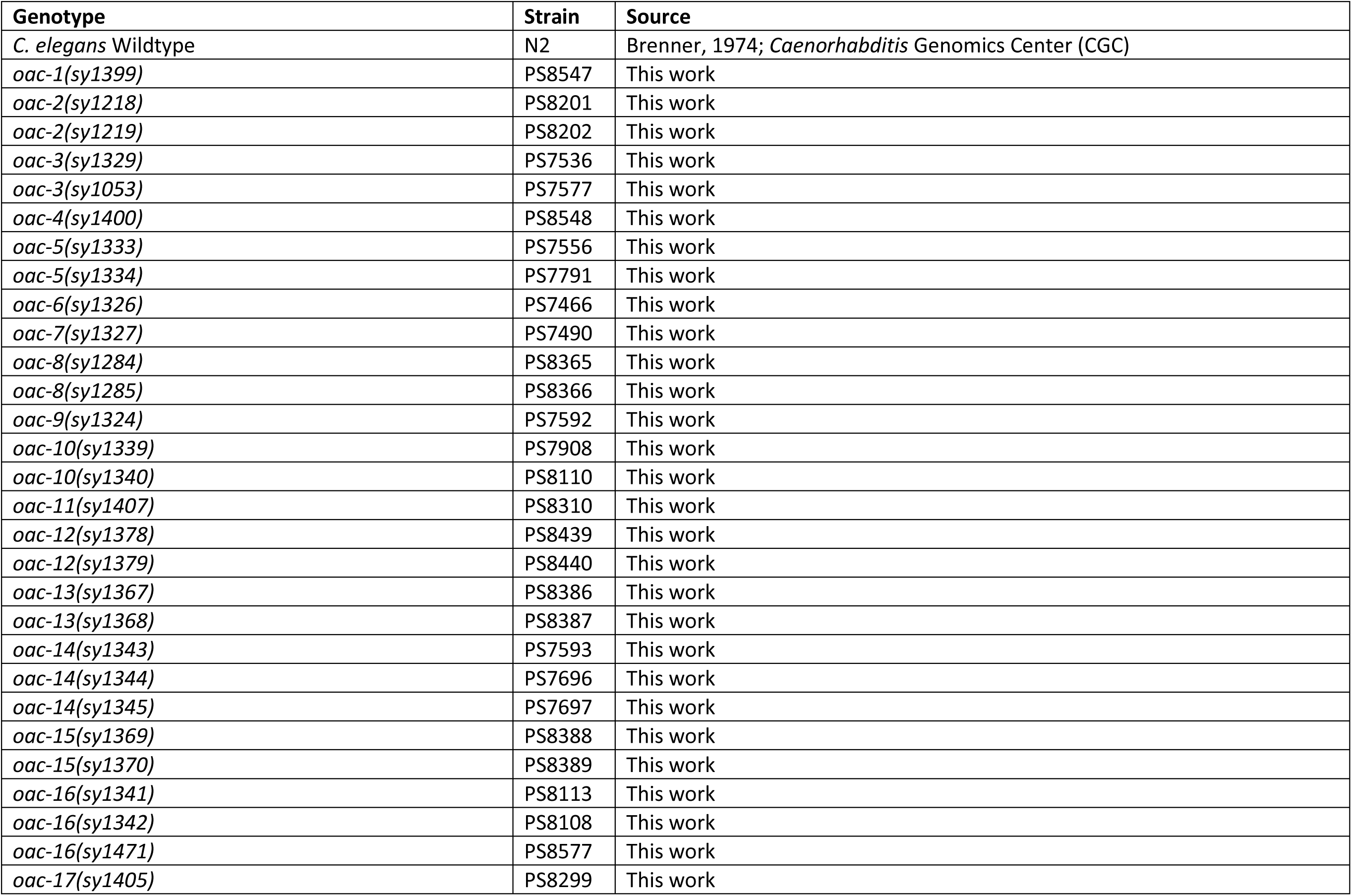
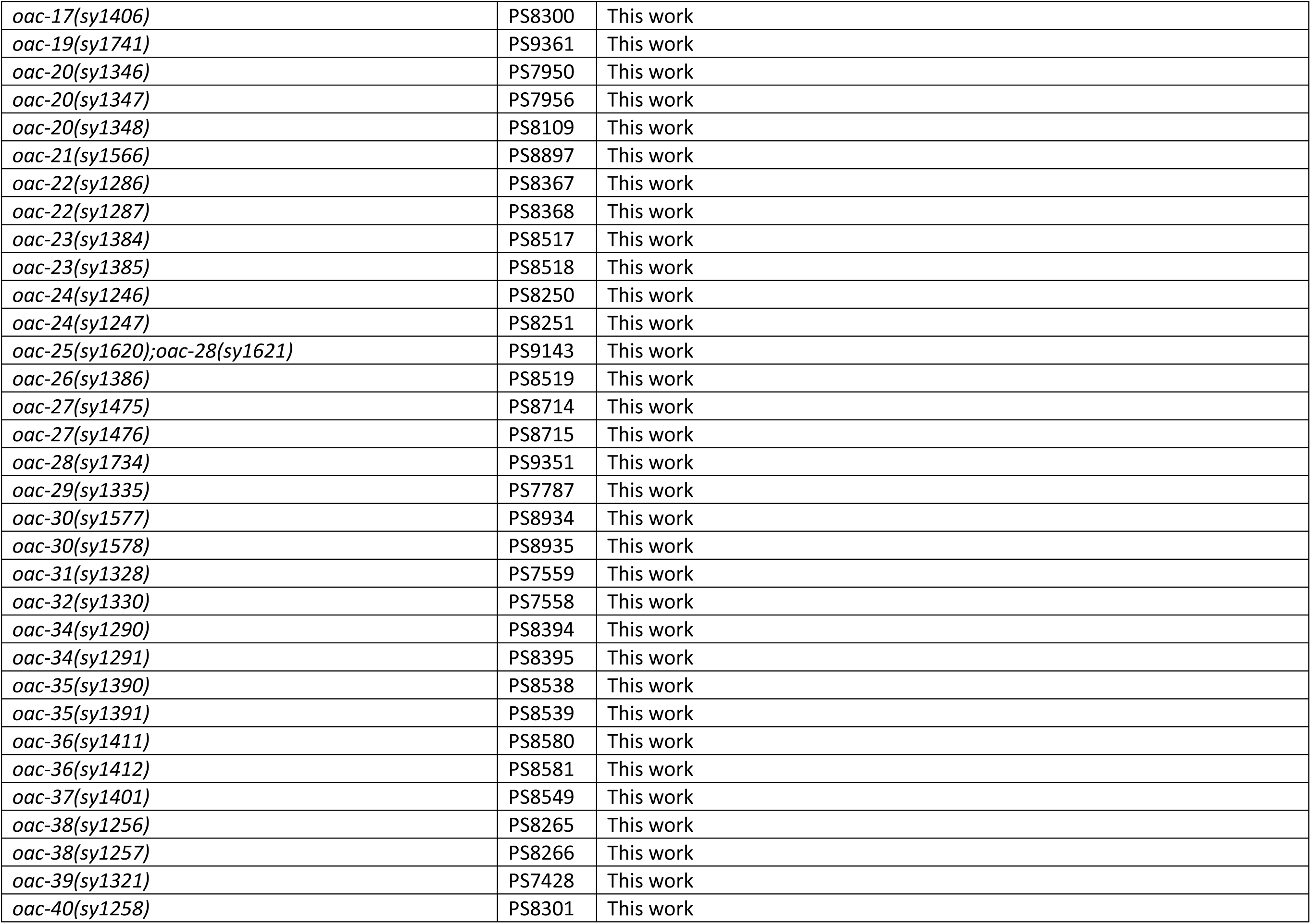
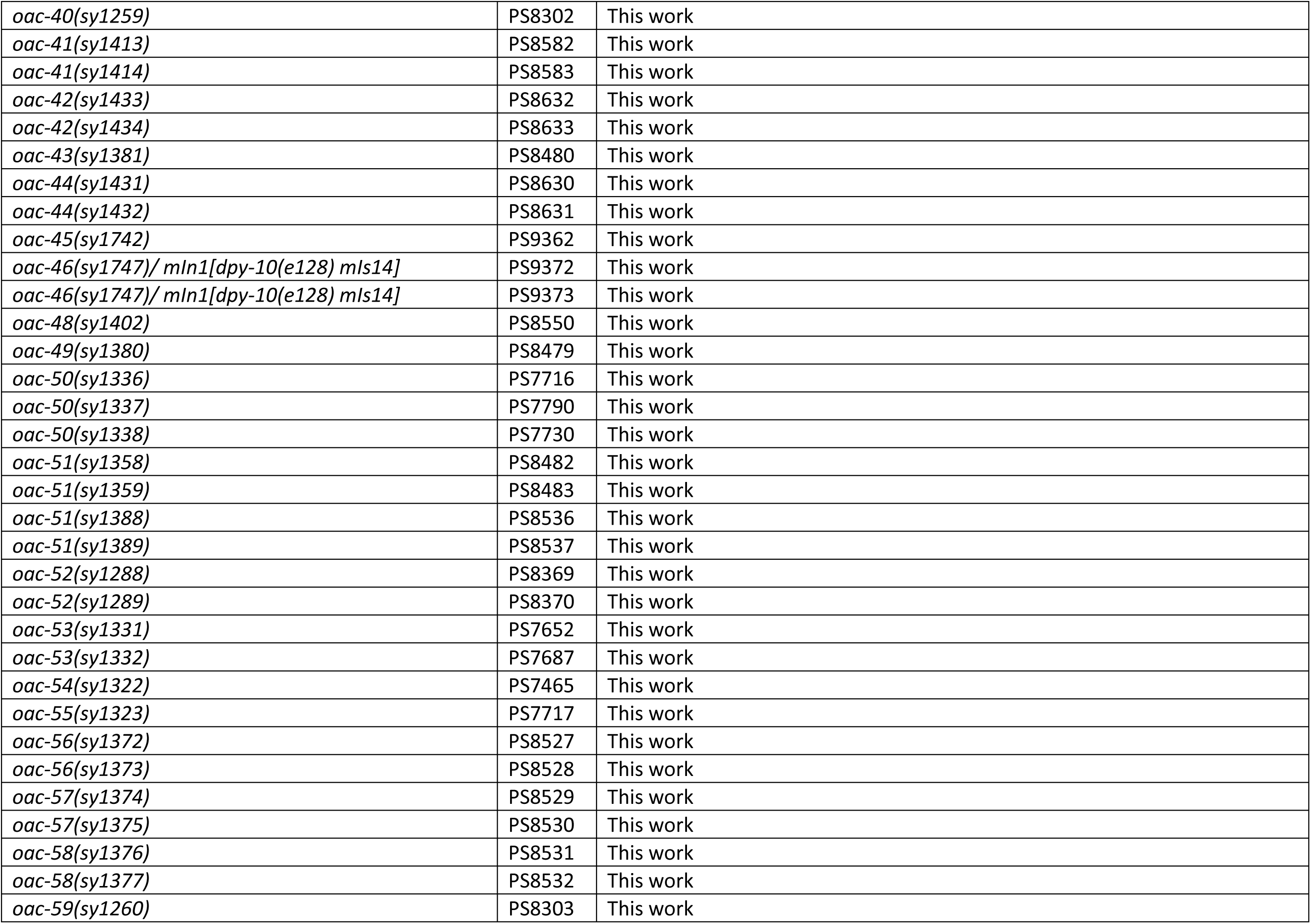
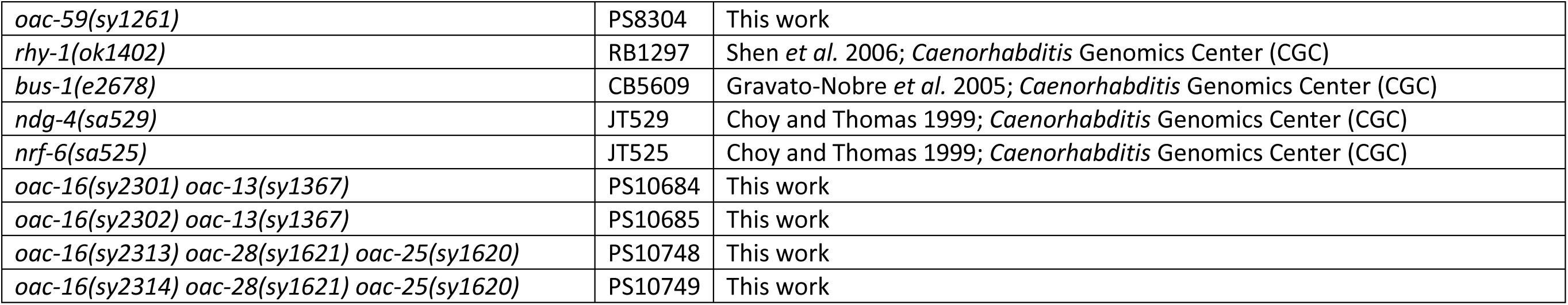
List of all *oac* mutant strains used in this paper.

